# Selective ablation of thymic and peripheral Foxp3^+^ regulatory T cell development

**DOI:** 10.1101/2023.08.04.551974

**Authors:** Acelya Yilmazer, Dimitra Maria Zevla, Rikke Malmkvist, Carlos Alejandro Bello Rodríguez, Pablo Undurraga, Emre Kirgin, Marie Boernert, David Voehringer, Olivia Kershaw, Susan Schlenner, Karsten Kretschmer

## Abstract

Foxp3^+^ Treg cells of thymic (tTreg) and peripheral (pTreg) developmental origin are thought to synergistically act to ensure immune homeostasis, with self-reactive tTreg cells primarily constraining autoimmune responses. We exploited tTreg-specific GFP/Cre recombinase activity to selectively ablate either tTreg (ΔtTreg) or pTreg (ΔpTreg) cell development. In contrast to the tTreg cell behavior in ΔpTreg mice, pTreg cells with a highly activated suppressor phenotype replenished the Treg cell pool of C57BL/6.ΔtTreg mice, preventing early mortality and fatal autoimmunity. Even with advancing age, pTreg cells largely maintained immune tolerance in C57BL/6.ΔtTreg mice. However, only two generations of (C57BL/6>NOD) backcrossing precipitated severe disease lethality associated with a distinct, partially overlapping pattern of organ-specific autoimmunity. Genetic association studies defined a small set of autoimmune risk loci sufficient to unleash a particularly severe form of diabetes, including genes known to impinge on Treg cell biology. Thus, pTreg cells exhibit an unexpectedly high functional adaptability, emphasizing their importance as mediators of bystander effects to ensure self-tolerance.

**SUMMARY:** This study in complementary loss-of-function mouse models uncovers an unexpected functional plasticity of pTreg cells in constraining systemic autoimmune responses in the absence of tTreg cells and identifies tTreg cells as primary regulators of β-cell autoimmunity in type 1 diabetes.

## INTRODUCTION

The discovery of genetic *Foxp3* gene mutations as the culprit of the fatal autoimmune syndrome in the spontaneous *scurfy* mouse mutant (Chatila et al., 2000; Brunkow et al., 2001) and human IPEX patients (Bennett et al., 2001; Wildin et al., 2001) provided the basis for unraveling the key role of Foxp3^+^ regulatory T (Treg) cells in dominant immunological tolerance. Observations in *Foxp3* gene-targeted mice further corroborated Treg cell paucity as the primary cause of early death and multi-organ autoimmunity in Foxp3-deficient mice (Fontenot et al., 2003) but also revealed the peripheral accumulation of Treg cell-like ‘wanna-be’ CD4^+^ T cells with self-reactive specificities (Hsieh et al., 2006; Lin et al., 2007; Lahl et al., 2009) that contribute to the disease pathology (Kuczma et al., 2009; Wyss et al., 2016). Acute Foxp3^+^ Treg cell ablation recapitulated some, but not all aspects of the *scurfy* syndrome in non-autoimmune-prone mice (Kim et al., 2007; Lahl et al., 2007) and highlighted the continuous requirement of Treg cells to constrain organ-specific autoimmune responses in the spontaneous NOD mouse model of human type 1 diabetes (T1D) (Watts et al., 2021).

Since then, it has become clear that the physiologic Treg cell pool is developmentally heterogeneous (Sakaguchi et al., 2008; Lee et al., 2011; Shevach and Thornton, 2014), consisting of intrathymically (tTreg) and peripherally (pTreg) induced Treg cells that originate from distinct CD4^+^CD25^high^Foxp3^−^ precursor cells residing in thymus (Lio and Hsieh, 2008) and peripheral lymphoid tissues (Schallenberg et al., 2010), respectively. In the thymus, distinct CD4^+^CD8^−^ single-positive (CD4SP) precursor cells, which exhibit low levels of Foxp3 protein preceding the up-regulation of CD25 expression, further expand the mature tTreg cell repertoire (Owen et al., 2019). In early studies examining the functional specialization of Treg cell developmental subsets by adoptive transfer immunotherapy of newborn *scurfy* mice (Haribhai et al., 2011), total Foxp3^+^ Treg cells prevented disease lethality, but did not suppress chronic inflammation and autoimmunity, which required the provision of Foxp3-sufficient CD4^+^ T cells to facilitate the extrathymic conversion of initially Foxp3^−^ T cells into functional Foxp3^+^ Treg cells (Haribhai et al., 2011). According to the prevailing view, tTreg cells are primarily positively selected by self-antigens during intrathymic development and are functionally specialized to control immune homeostasis and autoimmune responses (Sakaguchi et al., 2008; Pohar et al., 2018). The tTreg cell compartment in the spleen (SPL) and lymph nodes (LNs) has also been proposed to harbor Foxp3^+^ST-2^+^ common precursors for tissue-type Treg cells (Delacher et al., 2020) that accumulate and perform homeostatic and regenerative functions in nonlymphoid tissues (Campbell and Rudensky, 2020), such as the visceral adipose tissue (Feuerer et al., 2009; Cipolletta et al., 2012). Consistent with tTreg cells as primary regulators of autoimmune responses, studies in mice with *Foxp3* gene-targeted deletion of conserved non-coding region 1 (CNS1) (Foxp3.CNS1^−/−^), which exhibit a significant, albeit incomplete block of pTreg cell development (Zheng et al., 2010), failed to reveal severe autoimmune symptoms and have implicated pTreg cells in the control of immune responses at mucosal surfaces (Josefowicz et al., 2012) and maternal-fetal tolerance (Samstein et al., 2012). More recently, pTreg cells dependent on the gut microbiota have been shown to mediate functions beyond dominant suppression by facilitating muscle regeneration (Hanna et al., 2023). With regard to a putative role of pTreg cells in the control of autoimmune responses, previous studies in the NOD model showed that dendritic cell (DC)-targeted self-antigen can encourage highly diabetogenic CD4^+^Foxp3^−^ T cells to acquire a Foxp3^+^ pTreg cell phenotype (Kretschmer et al., 2005; Petzold et al., 2012), and that naturally induced, β cell-reactive pTreg cells are superior to tTreg cells with the same T cell receptor (TCR) specificity in constraining the manifestation of overt diabetes in a NOD.*Rag1*^−/−^ adoptive transfer model (Petzold et al., 2014). While Foxp3-deficient NOD mice failed to develop insulitis and overt diabetes (Chen et al., 2005), studies in Foxp3.CNS1^−/−^ NOD mice have provided ambiguous results, providing evidence for either a dispensable (Holohan et al., 2019) or nonredundant function (Schuster et al., 2018) of Foxp3.CNS1-dependent pTreg cells in the control of destructive β cell autoimmunity. In these studies, the relative contribution of tTreg cells to autoimmune β cell protection has not been directly addressed, owing to the lack of mouse models with selective tTreg cell paucity.

Here, we have exploited tTreg cell lineage-specific GFP/Cre recombinase activity in dual Foxp3^RFP/GFP^ reporter mice (Schallenberg et al., 2012; Petzold et al., 2014) to generate complementary mouse lines that are deficient in either the tTreg (Simonetti et al., 2023) or pTreg cell lineage, while sparing the respective sister population. The results of subsequent loss-of-function studies revealed an unexpectedly high functional adaptability of naturally occurring pTreg cells in mice with selective tTreg cell paucity, thereby preventing the manifestation of severe *scurfy*-like symptoms commonly observed in mice with complete Treg cell deficiency. However, the acquisition of an increased genetic autoimmune risk associated with compromised Treg cell activity unleashed high mortality and a distinct pattern of autoimmune diseases, including severe β cell autoimmunity and overt diabetes.

## MATERIALS and METHODS

### Selective *in vivo* Ablation of Developmental Treg Cell Sublineages

Foxp3^RFP/GFP^ mice (**Fig. 1A**) (Petzold et al., 2014), congenic CD45.1 *scurfy* mice, and Rag2^−/−^ mice were on the C57BL/6 (B6) background. NOD.Foxp3^RFP/GFP^ mice were obtained by backcrossing B6.Foxp3^RFP/GFP^ mice onto the NOD/ShiLtJ background (Jackson Laboratories, Bar Harbor, USA) for ≥14 generations (Petzold et al., 2014). For tTreg cell ablation (**Fig. 1B**), B6.R26^DTA^ mice with Cre-activatable diphtheria toxin A (DTA) expression from the ubiquitous *Rosa26* gene locus (Voehringer et al., 2008) were crossed with B6.Foxp3^RFP/GFP^ mice (Petzold et al., 2014; Simonetti et al., 2023) or backcrossed to NOD.Foxp3^RFP/GFP^ mice, as indicated. For pTreg cell ablation (**Fig. 1C**), a conditional Foxp3-STOP allele with Cre-activatable Foxp3 expression was crossed to B6.Foxp3^RFP/GFP^ mice, or backcrossed to NOD.Foxp3^RFP/GFP^ mice, as indicated. The Foxp3-STOP allele was developed at the University of Leuven (Genome Engineering Platform). In brief, a transcriptional STOP cassette, consisting of two loxP-flanked SV40 polyadenylation sites, was introduced by conventional gene targeting in E14 ES cells between exon 4 and 5 of the *Foxp3* gene, followed by excision of an FRT-flanked neomycin resistance gene using deleter mice [*Gt(ROSA)26Sor^tm1(FLP1)Dym^*/J; Stock No. 003946] (Farley et al., 2000). The conditional Foxp3-STOP allele was then backcrossed onto the B6 background for ≥ 10 generations and crossed with B6.Rag2^−/−^ mice, protecting B6.Foxp3-STOP mice from severe autoimmunity due to complete Foxp3^+^ Treg cell deficiency. All NOD mouse lines were fed with NIH #31M rodent diet (Altromin, Germany), and their blood glucose levels were routinely determined once a week using whole blood from the tail vein and Accu-Chek® Aviva (Roche). Mice were considered diabetic at blood glucose levels above 200 mg/dl on at least two consecutive measurements or with blood glucose levels once above 400 mg/dl. All mice were housed and bred at the Animal Facility of the CRTD under specific pathogen-free conditions. Animal experiments were performed as approved by the Landesdirektion Dresden (25-5131/502/5, TVA 5/2020; 25-5131/522/43, TVV41/2021).

**Figure 1:**
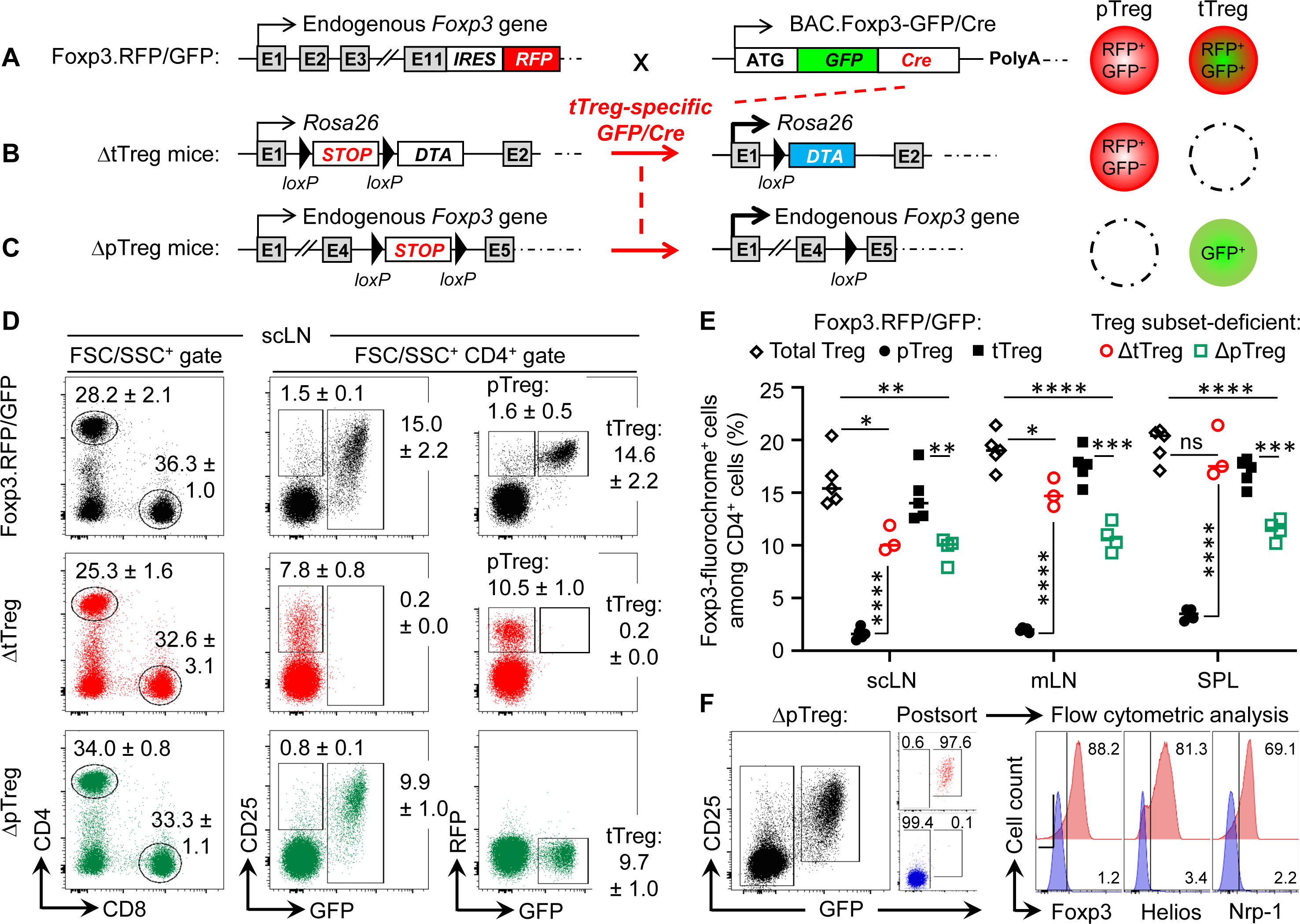
Selective ablation of tTreg and pTreg cell development *in vivo*. **(A-C)** Schematic overview of genetic strategy. **(A)** Foxp3^RFP/GFP^ mice. RFP is expressed from an IRES downstream of the endogenous *Foxp3* gene in both pTreg and tTreg cells. Restricted activation of BAC.Foxp3^GFP/Cre^ reporter expression to the thymus results in tTreg cell lineage-specific GFP/Cre activity and induction of gene expression by loxP-flanked STOP cassette excision. **(B)** ΔtTreg mice. Ablation of tTreg cells by GFP/Cre-mediated induction of diphtheria toxin A (DTA) expression. **(C)** ΔpTreg mice. A Foxp3.STOP cassette precludes pTreg cell development, while tTreg cell development can proceed after GFP/Cre-mediated induction of endogenous *Foxp3* gene expression. For this, the Foxp3^IRES-RFP^ reporter in **(A)** was replaced by a Cre-activatable Foxp3.STOP cassette. **(D-F)** Flow cytometry of Treg cells in peripheral lymphoid tissues. **(D)** Representative dot plots of (left) FSC/SSC-gated and (middle, right) CD4-gated cells from subcutaneous lymph nodes (scLNs) of 21-22-week-old males of indicated mouse lines. Numbers in dot plots represent mean percentages of cells ± SD within the respective gate. **(E)** Percentages of CD4-gated Foxp3-fluorochrome^+^ Treg cells in scLNs, mesenteric LNs (mLNs), and spleen (SPL) of Foxp3^RFP/GFP^ mice (pTreg: filled black circles, n = 5; tTreg: filled black squares, n = 5), ΔtTreg mice (pTreg: open red circles, n = 3), and ΔpTreg mice (tTreg: open green squares, n = 4). Symbols and horizontal lines represent individual mice and mean values, respectively. Unpaired t-test: ns, not significant; *p ≤ 0.05, **p ≤ 0.01, ***p ≤ 0.001, **** p ≤ 0.0001. **(F)** CD4^+^GFP^−^ T cells and CD4^+^CD25^+^GFP^+^ tTreg cells were FACS-purified from peripheral lymphoid organs of 3-5 males at 10 weeks of age and subjected to flow cytometric analysis of Foxp3, Helios, and Nrp-1 expression after intracellular staining using fluorochrome-conjugated mAbs. Numbers in dot plots and histograms represent the percentage of cells within the respective gate.

### Histopathology

After euthanizing the mice using CO_2_ inhalation, organs were collected and briefly washed in PBS. Subsequently, the tissues were fixed in a 4% paraformaldehyde solution (Sigma-Aldrich), paraffin-embedded, and 5 µm sections were cut. These sections were then stained with hematoxylin and eosin to assess histopathological changes. A total of 13 organs were examined (lung, heart, thymus, thyroid gland, stomach, liver, intestine, kidney, pancreas, urinary bladder, mesenteric adipose tissue, reproductive tract, brain) in a blinded manner to evaluate the presence and extent of inflammation and necrosis (none: 0; mild: +; moderate: ++; severe: +++) as described elsewhere (Telieps et al., 2013).

### Flow Cytometry and Cell sorting

All single cell suspensions were prepared in Hank’s buffer (1×HBSS, 5% FCS, 10mM HEPES; all ThermoFisher, Life Technologies). For this, thymus (THY), spleen (SPL), mesenteric lymph nodes (mLN), pancreatic LN (pLN), and a pool of subcutaneous LN (scLN) (*Lnn. mandibularis, Lnn. cervicales superficiales, Lnn. axillares et cubiti, Lnn. inguinales superficiales,* and *Lnn. subiliaci*) were meshed through 70 μm cell strainers (BD Biosciences). Bone marrow (BM) cells were harvested from femurs and tibias by flushing mechanically dissociated bones or intact bone cavities with Hank’s buffer, followed by filtration through 70 μm cell strainers (BD). Single cell suspensions from SPL and BM were subjected to red blood cell lysis (erythrocyte lysis buffer, EL; Qiagen). Monoclonal antibodies (mAbs) to B220 (RA3-6B2), CD3ε (145-2C11), CD4 (RM4-5), CD8α (53-6.7), CD25 (PC61), CD62L (MEL-14), CD44 (IM7), CD45.1 (A20), CD45.2 (104), CD103 (M290), c-Kit (2B8), GITR (DTA-1), ICOS (7E.17G9), KLRG1 (2F1), PD-1 (29F.1A12), ST-2 (U29-93), IgD (11-26c), IgM (II/41), MHC class II (I-A^b^: M5/114.15.2; I-A^G7^: OX-6), Foxp3 (FJK-16s), Helios (22F6), IL-10 (JES5-16E3), IFN-γ (XMG1.2), IL-17a (eBio17B7), IL-2 (JES6-5H4), IL-4 (11B11), IL-5 (TRFK5), TNF (MP6-XT22), Fc receptor-blocking mAb against CD16/32 (93), and fluorochrome-conjugated streptavidin (BUV395, eFlour450, APC and PE-Cy7) were purchased from BD, eBioscience, or Biolegend. Abs to Nrp1 (polyclonal goat IgG-AF700) were purchased from R&D Systems. Intracellular expression of cytokines and transcription factors was analyzed using the respective fluorochrome-coupled mAbs in conjunction with either the BD Cytofix/Cytoperm kit (BD) or the Foxp3 staining buffer set (eBioscience) according to the manufacturer’s protocol. The numbers of viable cells were determined using propidium iodide and a MACSQuant (Miltenyi Biotec). Before cell sorting, cells were enriched for CD4^+^ or CD25^+^ cells using biotinylated mAbs directed against CD4 or CD25, respectively, streptavidin-conjugated microbeads, fluorochrome-conjugated streptavidin, and the AutoMACS Pro magnetic cell separation system (Miltenyi Biotec). Samples were stained with DRAQ7 (BioStatus) for dead cell exclusion, filtered through 40 µm cell strainers, and analyzed on a LSR Fortessa or sorted using a FACS Aria II or III (all BD). Data were analyzed using FlowJo software (Version 10.8.1, Tree Star Inc.).

### Adoptive T Cell Transfer

Single cell suspensions from pooled LNs and SPL of B6.Foxp3^RFP/GFP^ donors (CD45.2) were subjected to CD4-based magnetic bead enrichment, followed by FACS-based isolation of total CD4^+^ T cells (*i.e.*, including GFP^+^ tTreg cells) and CD4^+^GFP^−^ T cells (*i.e.*, depleted of GFP^+^ tTreg cells). 1 x 10^7^ cells were injected i.p. into ≤ 2-day-old congenic CD45.1 *scurfy* recipient mice.

### T Cell Culture

T cells were cultured in 96-well round-bottom plates (Greiner) at 37°C and 5% CO_2_ in 200 μl RPMI complete medium [RPMI 1680 medium supplemented with 1 mM Sodium pyruvate, 1 mM HEPES, 2 mM Glutamax, 100 U/ml Penicillin, 100 µg/ml Streptomycin, 100 µg/ml Gentamycin, 0.1 mM non-essential amino acids, 0.55 mM β-mercaptoethanol and 10% FCS (v/v); all ThermoFisher, Life Technologies]. Prior to the flow cytometric analysis of intracellular cytokines, single cell suspensions were stimulated for 4 h in RPMI complete medium, using 50 ng/ml phorbol 12-myristate 13-acetate (PMA) and 200 ng/ml ionomycin (Iono), in the presence of 20 μg/ml Brefeldin A (all Merck, Sigma-Aldrich). For *in vitro* suppression, CD4^+^CD62L^high^CD25^−^Foxp3^−^ T responder (Tresp) cells and CD4^+^CD25^+^Foxp3^+^ Treg cells (RFP^+^GFP^+^ tTreg or RFP^+^GFP^−^ pTreg cells from B6.Foxp3^RFP/GFP^ mice; and GFP^+^ tTreg and RFP^+^GFP^−^ pTreg cells from B6 mice with selective pTreg and tTreg cell paucity, respectively) were FACS-isolated from peripheral lymphoid tissues. 5 × 10^4^ eFluor670-labeled (5 µM, eBioscience) Tresp cells were cultured in triplicate wells with 10^5^ T cell-depleted splenocytes (magnetic bead depletion using anti-CD3, -CD4, and -CD8 mAb) and soluble anti-CD3ε mAb (0.5 μg/ml), either alone or with Treg cells at different Treg:Tresp cell ratios, as indicated. On day 3 after initiation of cultures, Tresp cell proliferation and CD25 expression was assessed by flow cytometry.

### Genomic PCR-based *Idd* gene analysis

Genomic DNA was isolated from tail biopsies using the NucleoSpin DNA RapidLyse kit (Macherey-Nagel) according to the manufacturer’s protocol. Genomic PCR was performed using DreamTag green DNA polymerase and buffer, dNTPs (all Thermo Fisher, Life Technologies), a set of 36 primer pairs (Eurofins Genomics) that cover most of the known *Idd* loci (see **Supplementary Materials and Methods** for a complete list) (Serreze et al., 1996), and a Biometra Trio Thermocycler (Analytik Jena).

### Statistical analysis

Statistical significance was assessed using Prism 8 software (Version 8.4.3, GraphPad Software Inc., CA, USA). As indicated, the Student’s *t*-*test* (unpaired, two-tailed), Long-rank test (multiple comparisons with Bonferroni correction), and Chi-square test was used to assess statistical significance. Differences were considered as significant when *p ≤ 0.05, **p ≤ 0.01, ***p ≤ 0.001, ****p ≤ 0.0001.

## RESULTS

### tTreg-specific Cre activity enables selective blockage of tTreg and pTreg cell development

In dual Foxp3^RFP/GFP^ reporter mice (**Fig. 1A**), expression of the GFP/Cre fusion protein closely correlates with Foxp3 protein expression in the tTreg cell lineage, as transcriptional activation of the BAC.Foxp3^GFP/Cre^ reporter is restricted to the thymic *in vivo* environment (Schallenberg et al., 2012; Petzold et al., 2014; Junius et al., 2021; Simonetti et al., 2023). In brief, the BAC.Foxp3^GFP/Cre^ reporter was completely inactive in physiologic Foxp3^−^CD4^+^CD25^+^ pTreg cell precursors at peripheral sites (Schallenberg et al., 2010; Petzold et al., 2014), in various experimental settings of pTreg cell induction *in vivo*, and upon artificial Foxp3 induction in naïve CD4^+^Foxp3^−^ T cells *in vitro* (Petzold et al., 2014). Accordingly, thymic Foxp3^−^CD25^+^ CD4SP tTreg cell precursors upregulated Foxp3-driven GFP/Cre expression during developmental progression *in situ*, or after intrathymic injection, but not *in vitro* in IL-2-supplemented cultures (Petzold et al., 2014).

In Foxp3^RFP/GFP^ x R26^DTA^ mice (hereafter referred to as ΔtTreg mice) (Simonetti et al., 2023), GFP/Cre recombinase activity induces DTA expression selectively in the tTreg cell lineage by excision of an upstream loxP-flanked STOP cassette (Voehringer et al., 2008) (**Fig. 1B**). DTA-mediated ablation (≥ 99.8%) resulted in the absence of RFP^+^GFP^+^ tTreg cells in subcutaneous LNs (scLN) (**Fig. 1D**) and at other peripheral sites, such as mesenteric LN and SPL (**Fig. 1E**). A similarly high ablation efficiency was observed in peripheral lymphoid tissues of ΔtTreg mice with heterozygous (B6.R26^DTA^) and homozygous (B6.R26^DTA/DTA^) expression of the R26.STOP-DTA transgene, and over a wide age range (3-80 weeks; data not shown). As compared to Foxp3^RFP/GFP^ mice, selective tTreg cell paucity had no appreciable impact on the proportional distribution of CD4^+^ and CD8^+^ T cells in scLNs (**Fig. 1D**, left), but was accompanied by a > 6-fold increase in the percentage of RFP^+^GFP^−^ pTreg cells among CD4-gated T cells (Foxp3^RFP/GFP^ mice: 1.6 ± 0.5%; ΔtTreg mice: 10.5 ± 1.0%) (**Fig. 1D,E**). This marked increase in the pTreg cell population size of ΔtTreg mice largely compensated for the numerical impairment of the overall Treg cell pool in peripheral lymphoid tissues (**Fig.1E**, **Supplementary Fig. S1A**), but to a significantly lesser extent in peripheral blood (**Supplementary Fig. S1B**). In aged, > 1-year-old ΔtTreg mice, pTreg cells even exceeded total Treg cell frequencies of age-matched Foxp3^RFP/GFP^ mice (**Supplementary Fig. S1C**).

To obtain mice with selective pTreg cell paucity (hereafter referred to as ΔpTreg mice), the Foxp3^IRES-RFP^ reporter of Foxp3^RFP/GFP^ mice was replaced by breeding with a Cre-activatable Foxp3.STOP cassette (**Fig. 1C**). Hemizygous and homozygous Foxp3.STOP mice succumb to severe *scurfy* disease due to the absence of functional Foxp3 protein and Treg cells (Gerbaux et al., 2023). In ΔpTreg mice, BAC.Foxp3^GFP/Cre^-mediated activation of Foxp3 expression selectively in tTreg cell lineage-committed thymocytes allowed for the formation of a robust peripheral GFP^+^ tTreg cell compartment (**Fig. 1D,E**), while the extrathymic generation of pTreg cells remained precluded by the Foxp3.STOP cassette. Consistently, in peripheral lymphoid tissues of ΔpTreg mice, the expression of Foxp3, Helios, and Nrp-1 was absent in FACS-purified CD4^+^GFP^−^ cells, but readily detectable in CD4^+^GFP^+^ tTreg cells (**Fig. 1F**). Interestingly, selective pTreg cell deficiency resulted in a sustained reduction of the peripheral tTreg cell pool in adult (**Fig. 1E, Supplementary Fig. S1A**) and aged (**Supplementary Fig. S1C**) ΔpTreg mice, consistent with previous observations in Foxp3.CNS1^−/−^ mice with impaired pTreg cell development (Zheng et al., 2010; Josefowicz et al., 2012). These data, in conjunction with the ability of pTreg cells to compensate for the tTreg cell loss in ΔtTreg mice, suggest that pTreg cells are subject to less stringent constraints of the T cell receptor (TCR)-dependent clonal niche in peripheral lymphoid tissues, as compared to tTreg cells (Bautista et al., 2009; Leung et al., 2009; Moran et al., 2011).

### Lymphopoiesis in ΔpTreg and ΔtTreg mice

Acute and chronic inflammatory immune responses are well-known to modulate hematopoietic activity, as exemplified by the manifestation of severe lympho-hematopoietic defects during ontogeny of *scurfy* mice (Chen et al., 2010; Leonardo et al., 2010; Chang et al., 2012; Riewaldt et al., 2012). In the *scurfy* model, autoimmune-mediated thymic aberrations include severe post-developmental atrophy associated with enhanced apoptosis of CD4^+^CD8^+^ double-positive (DP) cells and concomitantly increased frequencies of CD4SP and CD8^+^ SP (CD8SP) cells (Riewaldt et al., 2012). Unexpectedly, the proportional distribution of DP and SP cells in young (3-4-week-old; data not shown) and adult (13-22-week-old) (**Fig. 2A**) ΔtTreg mice did not significantly differ from age-matched cohorts of ΔpTreg mice and Treg cell-proficient Foxp3^RFP/GFP^ mice. The average thymus size of adult ΔtTreg mice was only moderately reduced (< 2-fold; **Fig. 2B,C**). We occasionally observed rare cases (≤ 10%) of severe thymic atrophy, which were restricted to individual ΔtTreg mice (**Fig. 2B,C**) that additionally exhibited a markedly reduced body weight (ΔtTreg: mouse #1, 11.4 g; mouse #2-5, 28.4 ± 2.8 g; WT: 27.9 ± 1.4 g; ΔpTreg: 31.6 ± 4.1 g), while high frequencies of pTreg cells in peripheral lymphoid tissues remained unaffected (data not shown).

**Figure 2:**
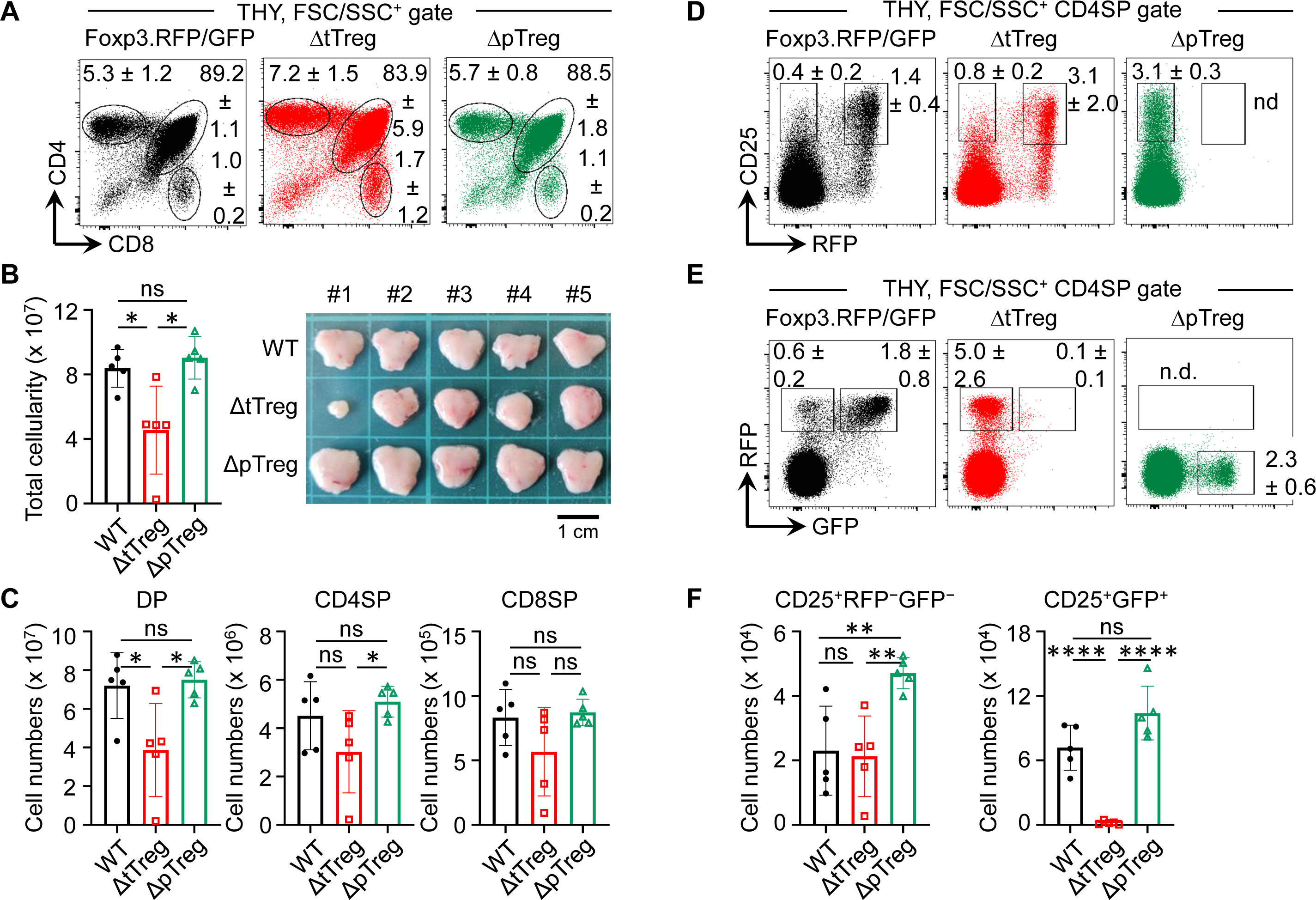
Thymopoiesis in adult ΔtTreg and ΔpTreg mice. Flow cytometry of thymic T cell development in Foxp3^RFP/GFP^, ΔtTreg, and ΔpTreg mice. **(A-C)** T cell development. **(A)** Representative flow cytometry of CD4 and CD8 expression among FSC/SSC-gated cells from the thymus (THY) of 17-22-week-old males, as indicated. **(B)** Total thymic cellularity (left) and thymus size (right). **(C)** Numbers of DP, CD4SP, and CD8SP cells. **(D-F)** tTreg cell development. Representative flow cytometry of **(D)** CD25 and Foxp3^IRES^-driven RFP expression, and **(E)** RFP and BAC.Foxp3^GFP/Cre^-driven GFP expression among gated CD4SP cells, as depicted in **(A)**. **(F)** Numbers of CD25^+^RFP^⎻^GFP^⎻^ (left) and CD25^+^GFP^+^ cells. Note that ΔpTreg mice lack the Foxp3^IRES-RFP^ reporter (n.d., not detectable). Numbers in dot plots in **(A,D,E)** represent mean percentages of cells ± SD within the respective gate. Symbols and bars in **(B,C,F)** represent individual mice and mean values ± SD, respectively. Unpaired t-test: ns, not significant; *p ≤ 0.05, **p ≤ 0.01, ****p ≤ 0.0001. Data are from a single experiment (5 mice per group) representative of 4 experiments performed (3-6 mice per experiment).

Next, we extended our observation of normal T lymphopoiesis in the majority of ΔtTreg mice to the analysis of B lymphopoiesis in the adult bone marrow (BM). In BM of *scurfy* mice, T cell-mediated autoimmune responses and systemically elevated levels of inflammatory cytokines have been shown to cause a complete block of B cell development, which is reflected by the absence of early Pro/Pre-B-I cell precursors and newly formed IgM^+^ B cells (Chen et al., 2010; Leonardo et al., 2010; Chang et al., 2012; Riewaldt et al., 2012). However, our comparative flow cytometric analyses failed to reveal evidence for dysregulated B lymphopoiesis in ΔtTreg and ΔpTreg mice, as compared to Treg cell-proficient Foxp3^RFP/GFP^ mice (**Supplementary Fig. S2**). This included comparable BM and spleen cellularity (**Supplementary Fig. S2A**), as well as proportions and numbers of early B220^+^c-kit^+^ Pro/Pre-B-I precursor cells and immature B220^low^IgM^+^ B cells in BM (**Supplementary Fig. S2B,C**), and of newly formed IgD^low^IgM^high^ B cells in the spleen (**Supplementary Fig. S2D,E**).

### DTA-mediated tTreg cell ablation occurs prior to thymic exit

In the thymus of Foxp3^RFP/GFP^ reporter mice, tTreg cell lineage commitment induces the sequential expression of RFP and GFP in initially Foxp3^−^CD25^+^ CD4SP cells (Petzold et al., 2014; Simonetti et al., 2023). Specifically, the developmental progression of Foxp3^−^CD25^+^ CD4SP cells first initiates the simultaneous up-regulation of Foxp3 and RFP protein (giving rise to CD25^+^RFP^+^GFP^−^ cells) (**Fig. 2D**; left panel), which is then followed by the timely delayed up-regulation of GFP/Cre expression, giving rise to newly formed Foxp3^+^CD25^+^ tTreg cells that are RFP^+^GFP^+^ (**Fig. 2E**; left panel). In ΔpTreg mice, flow cytometry revealed no adverse effects of selective pTreg cell paucity on tTreg cell development (**Fig. 2D,E**; right panels) and numbers of newly formed Foxp3^+^ tTreg cells (**Fig. 2F**). In the thymus of ΔtTreg mice, tTreg cell development proceeded to the CD25^+^RFP^+^GFP^−^ CD4SP stage (**Fig. 2D**; middle panel), but subsequent up-regulation of BAC.Foxp3^GFP/Cre^ reporter expression promoted DTA-mediated induction of apoptosis prior to thymic exit of newly formed CD25^+^RFP^+^GFP^+^ tTreg cells (**Fig. 2E**; middle panel; **Fig. 2F**). Thus, the observed deficiency in RFP^+^GFP^+^ tTreg cells at peripheral sites (**Fig. 1D,E**) was already established within the thymus. Whereas, the proportional increase of CD4-gated RFP^+^GFP^−^ cells in ΔtTreg mice (**Fig. 2D,E**) could be attributed to the intrathymic accumulation of mature pTreg cells (**Supplementary Figure S3A**) that originated from peripheral lymphoid tissues recirculating to the thymus, as indicated by heterogeneous CD25 expression levels and a ‘recirculating’ CD62L^−^CD69^+^CD44^high^ phenotype (**Supplementary Figure S3B**) (McCaughtry et al., 2007; Thiault et al., 2015).

### pTreg cells prevent the early manifestation of severe autoimmunity in ΔtTreg mice

In our B6 *scurfy* colony maintained under SPF conditions, approximately 50% of mice succumb to premature death by 35 days of age due to fatal autoimmunity associated with Foxp3 deficiency, and no mice live beyond 50 days (Riewaldt et al., 2012). Within 35 days after birth, ΔtTreg (**Fig. 3A**) and ΔpTreg (**Fig. 3B**) mice appeared overall healthy, showing no appreciable spontaneous mortality (**Fig. 3A,B**) or other *scurfy*-like symptoms (scaliness and crusting of eyelids/ears/tail, hepatomegaly, splenomegaly, lymphadenopathy, *etc.*) (data not shown). Consistently, our analysis of genotype distribution among 5-week-old mice revealed an unbiased heredity of the ΔtTreg (**Fig. 3C**) and ΔpTreg (**Fig. 3D**) phenotype.

**Figure 3:**
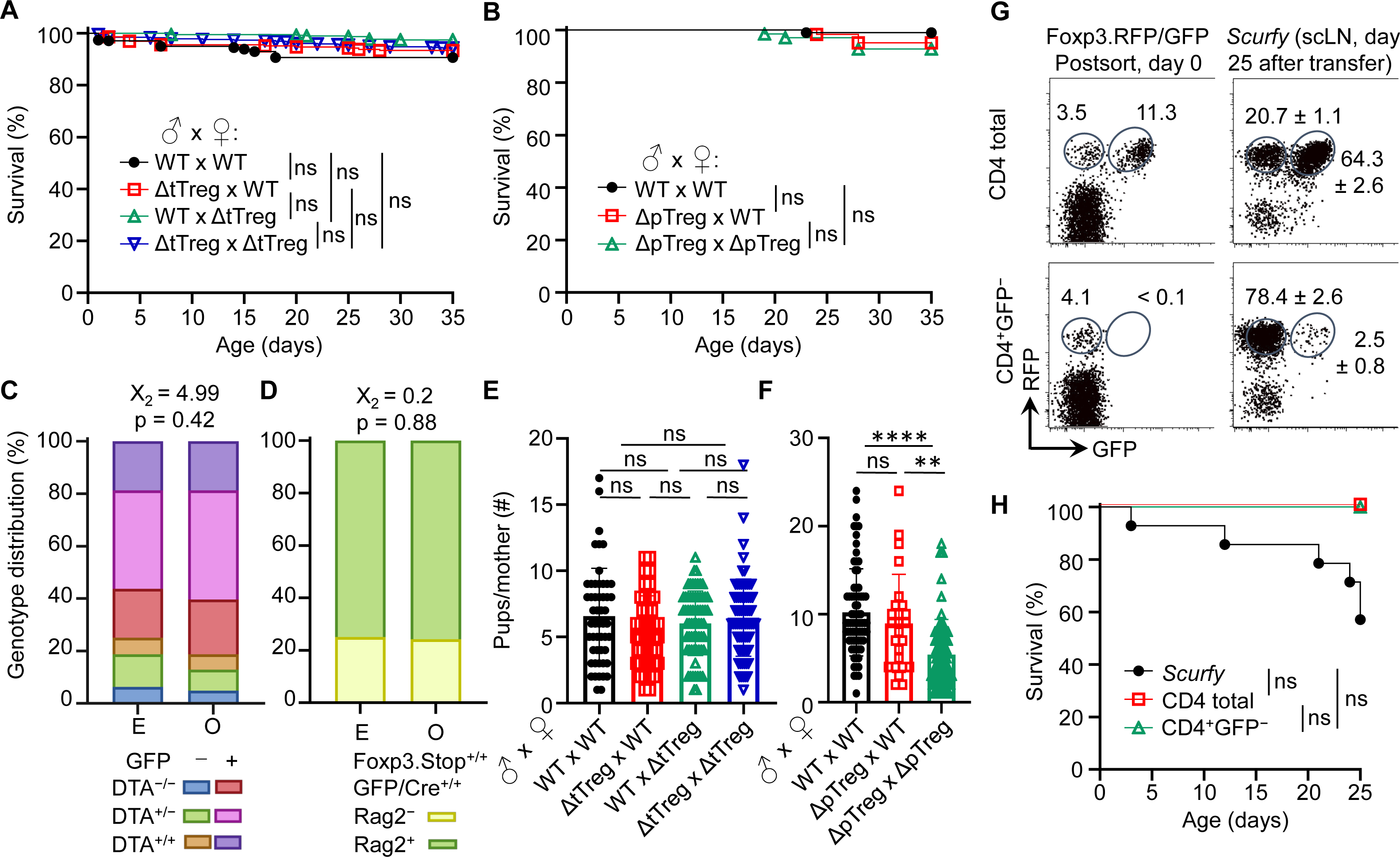
Viability and reproductive ability of ΔtTreg and ΔpTreg mice. **(A,B)** Kaplan-Meier survival curves of offspring produced by **(A)** ΔtTreg and **(B)** ΔpTreg pair mating. Newborn mice produced by indicated parental genotype combinations were monitored for the occurrence of spontaneous cases of mortality from birth onwards for up to 35 days (parental genotype, number of offspring; **(A)**: WT x WT, n = 342; ΔtTreg x WT, n = 227; WT x ΔtTreg, n = 398; ΔtTreg x ΔtTreg, n = 463. **(B)**: WT x WT, n = 103; ΔpTreg x WT, n = 62; ΔpTreg x ΔpTreg, n = 69). **(C,D)** Expected (E) and observed (O) distribution of offspring according to their genotype produced by **(C)** ΔtTreg (♂ DTA^+/−^GFP^+/−^ x ♀ DTA^+/−^GFP^+/−^; n = 187) and **(D)** ΔpTreg (♂ Rag2^+/−^Foxp3.Stop^+^GFP/Cre^+^ x ♀ Rag2^+/−^Foxp3.Stop^+/+^ GFP/Cre^+/+^; n = 62) pair mating. Chi-square test: X_2_, Chi-square. **(E,F)** The cumulative number of newborn pups produced by **(E)** ΔtTreg and **(F)** ΔpTreg pair mating of indicated parental genotype combinations. ΔtTreg: DTA^+/−^ or ^+/+^, BAC.Foxp3^GFP/Cre+^; ΔpTreg: Rag2^+/−^, Foxp3.Stop^hemi/homo^, BAC.Foxp3^GFP/Cre+^. Parental genotype, number of litters: **(E)**: WT x WT, n = 52; ΔtTreg x WT, n = 42; WT x ΔtTreg, n = 66; ΔtTreg x ΔtTreg, n = 72. **(F)**: WT x WT, n = 75; ΔpTreg x WT, n = 25; ΔpTreg x ΔpTreg, n = 60). Symbols and bars represent individual litters and mean values, respectively. **(G,H)** Adoptive Treg cell transfer into neonatal *scurfy* mice. **(G)** Flow cytometry of RFP and GFP expression among CD4-gated T cells before (post sort, left panels) and after (day 25, right panels) injection into conventional *scurfy* recipient mice. Numbers in dot plots represent percentages of cells (left) or mean percentages of cells ± SD (right) within the respective gate. **(H)** Kaplan-Meyer survival curves of *scurfy* mice that had either been left untreated (closed black circles, n = 14) or neonatally injected *i.p.* (5 x 10^5^ cells, day 0) with either total CD4^+^ T cells (open black squares, n = 4) or CD4^+^GFP^−^ T cells (open green triangles, n = 3) that had been FACS-purified from peripheral lymphoid tissues of Foxp3^RFP/GFP^ mice. At day 25, adoptively transferred CD4^+^CD45.2^+^ cells were tracked by flow cytometry in scLNs of congenic CD45.1^+^ recipient mice and analyzed for RFP and GFP expression. **(A,B,H)** Log-rank test and Bonferroni correction: ns, not significant. (**E,F**) Unpaired t-test: ns, not significant; **p ≤ 0.01, ****p ≤ 0.0001.

The impaired generation of pTreg cells in Foxp3.CNS1^−/−^ mice has been reported to impinge on maternal-fetal tolerance by increasing the resorption of semiallogeneic fetuses (Samstein et al., 2012). Our data show that the number of viable offspring produced by syngeneic ΔtTreg pair mating did not significantly differ from that of Foxp3^RFP/GFP^ mice (**Fig. 3E**). In contrast, interstrain breeding of ΔpTreg mice gave rise to significantly reduced numbers of offspring, correlating with the ΔpTreg phenotype of the breeding female (**Fig. 3F**). These findings in ΔtTreg and ΔpTreg mice, in conjunction with impaired implantation of syngeneic embryos after maternal Foxp3^+^ Treg cell depletion (Kahn and Baltimore, 2010; Heitmann et al., 2017), imply that pTreg cells may contribute to maternal-fetal tolerance even in syngeneic pregnancy.

The unexpected absence of severe *scurfy*-like symptoms in young B6.ΔtTreg mice suggested that selective tTreg cell paucity can be largely compensated by increased pTreg cell numbers (**Fig. 1D,E**). We next asked whether the observed pTreg cell behavior in the ΔtTreg model can be recapitulated in *scurfy* mice neonatally injected with CD4^+^GFP^−^ T cell populations (including RFP^+^GFP^−^ pTreg cells) that had been FACS-isolated from peripheral lymphoid tissues of Foxp3^RFP/GFP^ mice for RFP^+^GFP^+^ tTreg cell depletion (**Fig. 3G**, bottom left). In these experiments, total CD4^+^ T cells (including RFP^+^GFP^−^ pTreg and RFP^+^GFP^+^ tTreg cells) were included for comparison (**Fig. 3G**, top left). Both cohorts of *scurfy* recipients were viable (**Fig. 3H**) and appeared phenotypically healthy until the end of the observation period, apart from mild symptoms of delayed growth and exfoliative dermatitis in individual mice that received tTreg cell-depleted CD4^+^GFP^−^ T cells (data not shown). In the absence of tTreg cells, the adoptive CD4^+^GFP^−^ T cell transfer resulted in a marked accumulation of RFP^+^GFP^−^ pTreg cells among CD4^+^ T cells (day 0: 4.1%; day 25: 78.4± 2.6%) in *scurfy* recipients (**Fig. 3G**, bottom panels), most likely due to both the proliferative expansion of preformed pTreg cells and the conversion of initially CD4^+^Foxp3^−^ T cells (Haribhai et al., 2011). In *scurfy* recipients of total CD4^+^ T cells, RFP^+^GFP^−^ pTreg cell frequencies among CD4-gated cells also increased (day 0: 3.5%; day 25: 20.7 ± 1.1%), but the initial pTreg:tTreg cell ratio of 1:3 was preserved (**Fig. 3G**, top panels). Overall, these data indicate that pTreg cells can fill up the tTreg cell niche in both ΔtTreg (**Fig. 1D,E**) and *scurfy* mice (**Fig. 3G,H)** while maintaining a stable RFP^+^GFP^−^ phenotype.

### Maintenance of T cell homeostasis in adult ΔtTreg mice

Our initial characterization of young ΔtTreg mice failed to reveal evidence for disease symptoms associated with selective tTreg cell paucity. Consistently, mild immune infiltrations were limited to the salivary gland (mouse #1), and the lung (mouse #1) of individual ΔtTreg mice, but could not be observed in other organs, such as the liver or thyroid gland (**Fig. 4A**). Interestingly, although B6.ΔtTreg mice maintained normoglycemia, histological analyses consistently revealed pronounced immune infiltrates in the pancreas (**Fig. 4A**, bottom panels), which is in contrast to previous studies in other settings of tTreg cell deficiency, including Foxp3-deficient mice (Chen et al., 2005) and acute Treg cell ablation in DEREG mice on the B6 and NOD genetic background (Watts et al., 2021).

**Figure 4:**
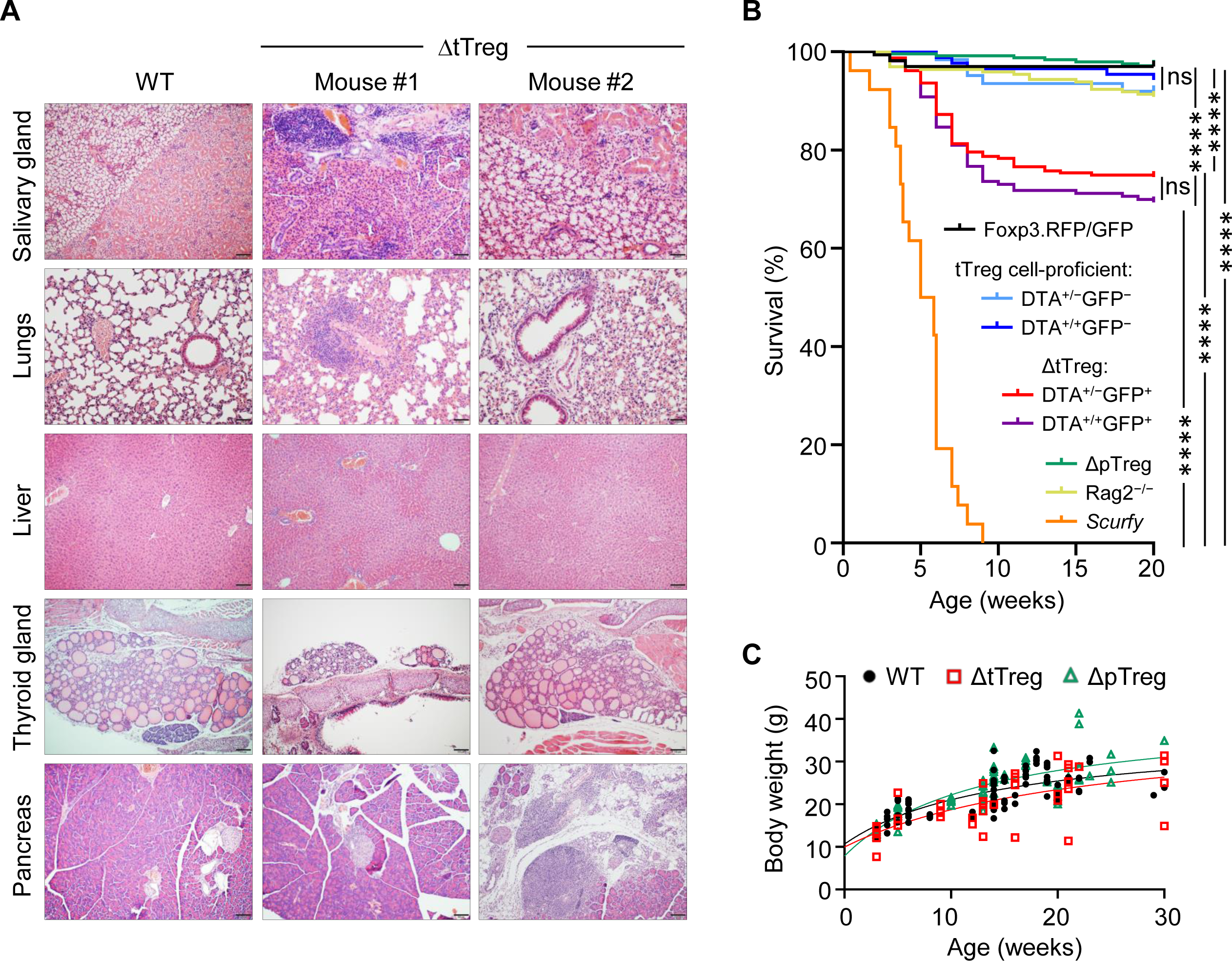
Autoimmune pathology and mortality of ΔtTreg mice. **(A)** Histological analysis. Organs of 4-week-old Foxp3^RFP/GFP^ (WT, left) and ΔtTreg (mouse #1 and #2) male mice were fixed in 4% PFA, and cut sections were stained with hematoxylin and eosin. In contrast to WT controls, individual ΔtTreg mice show mild leukocytic infiltrations in the salivary gland (mouse #1: +; mouse #2: ±), lungs (mouse #1: ++; mouse #2: ±), and pancreas (mouse #1: ++; mouse #2: ++). Sections from the liver and thyroid gland of WT and ΔtTreg mice lack leukocytic infiltrates and show normal organ structure. **(B)** Kaplan-Meier survival analysis. Cohorts of Foxp3^RFP/GFP^ mice (n = 167), ΔtTreg mice (DTA^+/−^GFP^+^, n = 235; DTA^+/+^GFP^+^, n = 163), and ΔpTreg mice (n = 245) were monitored for the occurrence of spontaneous cases of mortality from birth onwards for up to 20 weeks, as indicated. R26^DTA^ mice lacking the BAC.Foxp3^GFP/Cre^ transgene (DTA^+/−^GFP^−^, n = 62; DTA^+/+^GFP^−^, n = 87), Foxp3-deficient *scurfy* mice (n = 26), and immunodeficient Rag2^⎻/⎻^ mice (n = 196) were included for comparison. Log-rank test and Bonferroni correction: ns, not significant; ****p ≤ 0.0001. **(C)** Age-dependent body weight gain of Foxp3^RFP/GFP^ (WT, closed black circles, n = 92), ΔtTreg (open red squares, n = 38), and ΔpTreg (open red triangles, n = 60) mice.

With the advancing age of ΔtTreg mice, we noticed rare cases of spontaneous deaths, which became first apparent at the age of 7 weeks (R26^DTA^:18.7%, R26^DTA/DTA^: 19.0%; **Fig. 4B**). The mortality of ΔtTreg mice further increased thereafter, reaching a plateau by 10 weeks that was maintained until the end of the 20-week observation period (R26^DTA^:25.5%, R26^DTA/DTA^:30.7%), remaining well below the high mortality of *scurfy* mice (**Fig. 4B**). We rarely observed cases of spontaneous mortality among immunodeficient Rag2^−/−^ mice, or cohorts of tTreg cell-proficient, BAC.Foxp3^GFP/Cre–^ ΔtTreg littermates, ΔpTreg mice, and Foxp3^RFP/GFP^ mice (**Fig. 4B**). We further noticed that mortality appeared to be associated in part with a reduced body weight and thymic atrophy of the affected ΔtTreg mice (**Fig. 2B**). Our subsequent analyses showed that the majority of ΔtTreg mice had a body weight corresponding to their age (85.3%), but also confirmed that individual mice failed to keep up with physiological body weight gain (**Fig. 4C**), while the small and large intestine yielded an unsuspicious histopathological result (data not shown).

With regard to peripheral T cell homeostasis in adult ΔtTreg mice, total cellularity of scLNs (but not of mLNs and spleen; see also **Supplementary Fig. S2A**) was moderately, although significantly increased, as compared to Foxp3^RFP/GFP^ and ΔpTreg mice (**Fig. 5A**). We only occasionally observed adult ΔtTreg mice with pronounced lymphadenopathy and splenomegaly (< 10%; **Fig. 5A**). The analysis of inflammatory cytokine production indicated moderately increased proportions of IFN-γ- and IL-4-producing CD4^+^ T cells (**Fig. 5B**, top), and of IFN-γ-producing CD8^+^ T cells (**Fig. 5B**, bottom) in scLNs (but not in mLN or spleen; data not shown) of ΔtTreg mice. Increased expression of other cytokines (*e.g.*, IL-2, IL-10, IL-17) could also not be observed (data not shown). Consistent with largely normal frequencies of CD4^+^ and CD8^+^ T cells in the majority of ΔtTreg mice (**Fig. 1D**), our flow cytometric analysis of CD62L and CD44 expression revealed no evidence for systemically uncontrolled activation of CD4^+^ and CD8^+^ T effector cells, neither in scLNs (**Fig. 5C-E**) or other peripheral lymphoid tissues, such as mLN or spleen **(Fig. 5D,E)**.

**Figure 5:**
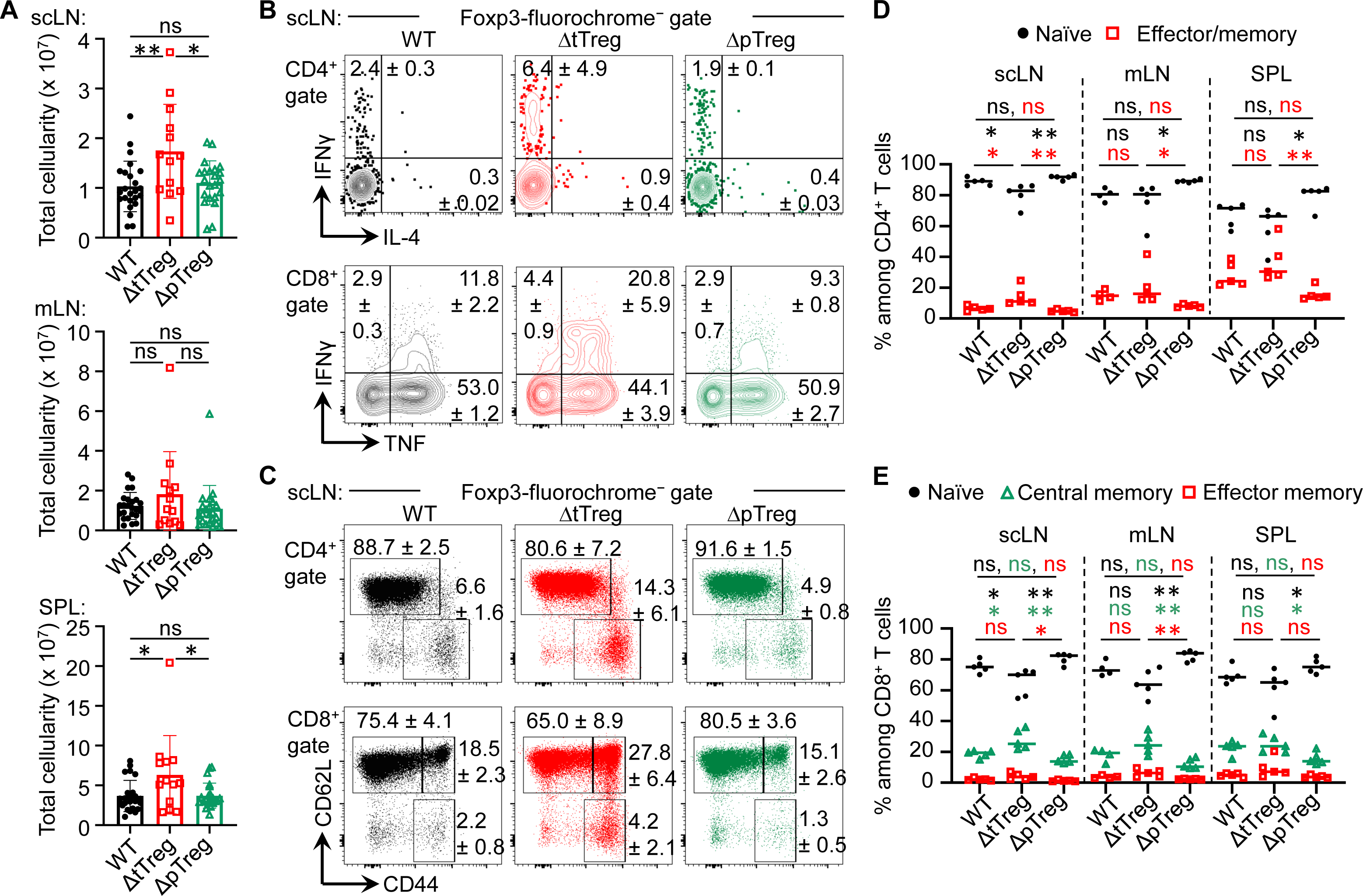
Peripheral T cell homeostasis in adult ΔtTreg and ΔpTreg mice. **(A)** Total cellularity of scLN (top), mLN, (middle), and SPL (bottom) of 13-22-week-old Foxp3^RFP/GFP^ (WT, n = 24), ΔtTreg (n = 13) and, ΔpTreg (n = 22) male mice. Symbols and bars represent individual mice and mean values ± SD, respectively. **(B-E)** Flow cytometry of CD4^+^ and CD8^+^ T effector cells in peripheral lymphoid tissues. **(B)** Representative dot plots of IFN-γ and IL-4 expression among gated CD4^+^Foxp3^−^ (top), and IFN-γ and IL-4 expression among gated CD8^+^ (bottom) T cells in scLN of 38-40-week-old WT (Foxp3^RFP/GFP^), ΔtTreg, and ΔpTreg male mice, as indicated (2-3 mice per group). **(C)** Representative dot plots of CD44 and CD62L expression among gated CD4^+^Foxp3^−^ (top) and CD8^+^ (bottom) T cells in scLN of 17-22-week-old WT (Foxp3^RFP/GFP^), ΔtTreg, and ΔpTreg male mice, as indicated (4-5 mice per group). (Gates: CD62L^high^CD44^low^, naïve; CD62L^low^CD44^high^, effector/memory; CD62L^high^CD44^high^, central memory). **(D,E)** Composite percentages of T cell effector subsets in 17-22-week-old Foxp3^RFP/GFP^ (WT), ΔtTreg, and ΔpTreg mice (4-5 male mice per group). **(D)** Naïve (CD62L^high^CD44^low^) and effector/memory (CD62L^low^CD44^high^) CD4^+^ T cell compartments. **(E)** Naïve (CD62L^high^CD44^low^), central memory (CD62L^high^CD44^high^), and effector/memory (CD62L^low^CD44^high^) CD8^+^ T cell compartments. Naïve: black closed circles; central memory: open green triangles; effector/memory: open red squares. Symbols and horizontal lines in **(A,D,E)** represent individual mice and mean percentages of cells ± SD, respectively. Numbers in dot plots in **(B,C)** indicate mean percentages of cells ± SD within the respective gate or quadrant. Unpaired t-test: ns, not significant; *p ≤ 0.05, **p ≤ 0.01.

### Compensatory adaptation of pTreg cell activity in the absence of tTreg cells

Despite its already high percentage share in steady-state Foxp3^RFP/GFP^ mice (Petzold et al., 2014), flow cytometric immunophenotyping indicated that the percentage of pTreg cells with a CD62L^low^CD44^high^ effector/memory-like phenotype further increased in LNs of adult ΔtTreg mice (**Fig. 6A**), which was in contrast to tTreg cells in ΔpTreg (**Fig. 6B**) mice. Consistent with an overall activated phenotype, pTreg cells in peripheral lymphoid tissues of ΔtTreg mice exhibited significantly upregulated expression of CD25 and several other ‘Treg cell signature’ proteins with functional relevance (including CD103, ICOS, ST2, and KLRG1), as compared to pTreg cells from Foxp3^RFP/GFP^ mice (**Fig. 6C**). In contrast, the reduced accumulation of pTreg in peripheral blood of ΔtTreg mice (**Supplementary Fig. S1B**) was accompanied by low levels of CD25 expression (**Fig. 6C**; left panel). Additionally, tTreg cells of ΔpTreg mice exhibited neither an increased effector/memory-like compartment (**Fig. 6B**) nor up-regulated ‘Treg cell signature’ protein expression (**Fig. 6C**), as compared to their tTreg cell counterparts in Foxp3^RFP/GFP^ mice. Consistently, pTreg cells isolated from ΔtTreg mice suppressed the activity of Tresp cells more efficiently in standard cocultures, as judged by the inhibition of Tresp cell proliferation and CD25 expression, and as compared with tTreg cells from ΔpTreg mice or total Treg cells from Foxp3^RFP/GFP^ mice (**Fig. 6D**). In summary, in contrast to tTreg cells of ΔpTreg mice, pTreg cells in peripheral lymphoid tissues (but not blood) of ΔtTreg mice acquire a highly activated state and increased suppressor function, which is indicative for their active involvement in constraining chronic immune dysregulation in the absence of tTreg cells **(Fig. 4)**.

**Figure 6:**
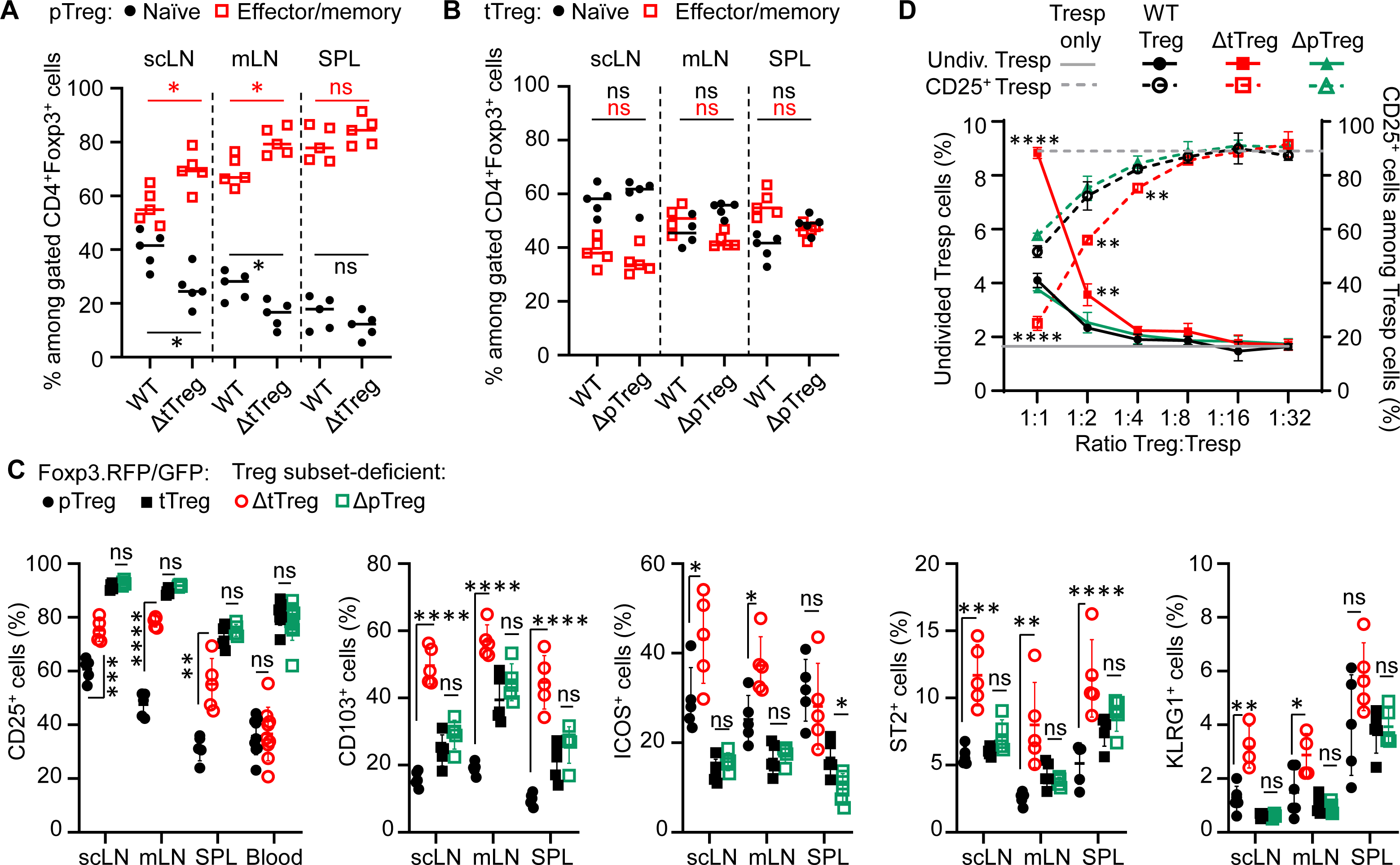
pTreg cell adaptation to tTreg cell paucity. **(A,B)** Flow cytometry of naïve and memory/effector-type Treg cell subsets. Cumulative percentages of **(A)** pTreg cells and **(B)** tTreg cells with a naïve (CD62L^high^CD44^low^, black filled circles) and effector/memory-type (CD62L^low^CD44^high^, red open squares) phenotype in peripheral lymphoid tissues (scLN, mLN, SPL) of ΔtTreg mice and ΔpTreg mice, respectively. Foxp3^RFP/GFP^ (WT) mice were included for comparison. Treg cell gating was as in Fig. 1D. **(C)** Signature protein expression. Composite percentages of surface marker expression among Foxp3-fluorochrome reporter-gated CD4^+^ Treg cells. Treg cell gating was as indicated in Fig. 1D: pTreg (closed black circles) and tTreg cells (closed black squares) of Foxp3^RFP/GFP^ mice; pTreg cells of ΔtTreg mice (open red circles); and tTreg cells of ΔpTreg mice (open green squares). Symbols and horizontal lines indicate individual mice and mean values ± SD, respectively. Data are from a single experiment, representative of 4 independent experiments performed (n = 3-6 per group; age: 13-22 weeks). **(D)** Suppressor function *in vitro*. Composite percentages of cell division (left, closed symbols) and CD25 expression (right, open symbols) of CD4^+^ Tresp cells at day 3 of co-culture, using total Treg cells of Foxp3^RFP/GFP^ mice (WT, black circles), pTreg cells of ΔtTreg mice (red squares), and tTreg cells (green triangles) of ΔpTreg mice at indicated Tresp:Treg ratios. For this, FACS-purified CD4^+^CD62L^+^Foxp3^−^CD25^−^ Tresp cells were co-cultured with APCs and 0.5 μg/ml α-CD3ε mAb, in the absence or presence of total Treg cells from scLN of Foxp3^RFP/GFP^ mice (WT, tTreg + pTreg), pTreg cells of ΔtTreg, or tTreg cells of ΔpTreg mice. Symbols and error bars in graphs indicate mean percentages ± SD of technical replicates (n = 2-3) from one experiment, representative of three independent experiments (5-10 mice per group at 20-22 weeks of age). Unpaired t-test: ns, not significant; *p ≤ 0.05, **p ≤ 0.01, ***p ≤ 0.001, ****p ≤ 0.0001.

### Fatal autoimmune pathology in ΔtTreg mice on a mixed (B6>NOD) background

Our characterization of ΔtTreg mice on the B6 background revealed neither early- or late-onset of severe morbidity nor other signs of fatal autoimmunity (**Fig. 3A, 4A,B**) consistently observed in mice with complete Foxp3^+^ Treg cell deficiency (Brunkow et al., 2001; Fontenot et al., 2003). However, the severity of autoimmune pathology associated with Treg cell deficiency can be markedly shaped by genetic factors: B6 *scurfy* mice can survive for up to 9 weeks after birth (**Fig. 4B**), which is significantly longer than *scurfy* mice on the BALB/c background, all of which succumb to death within < 5 weeks of birth (Haribhai et al., 2011). Here, we aimed to explore how increased genetic susceptibility impinges on the survival and immune homeostasis of ΔtTreg mice by backcrossing the B6.ΔtTreg mouse line to autoimmune-prone NOD mice carrying the dual Foxp3^RFP/GFP^ reporter. On a pure NOD background, spontaneous T1D manifestation is under polygenic control of more than 20 insulin-dependent diabetes (*Idd*) gene loci (Driver et al., 2011), which additionally confer a broad susceptibility to multiple other autoimmune syndromes (peripheral neuropathy, autoimmune thyroiditis, *etc.*), albeit often with a low incidence (Aubin et al., 2022). The analysis of (B6>NOD) hybrid mice obtained by two consecutive backcrosses (F2) revealed that selective tTreg cell paucity drastically decreased the survival of F2 ΔtTreg mice (both I-Ag7^+/–^ and I-Ag7^+/+^; see below) to ≤ 20% within 20 weeks after birth (**Fig. 7A**), as compared to F1 ΔtTreg mice (84.2%; **Fig. 7A**) and ΔtTreg mice on a pure B6 background (R26^DTA^: 74.5%, R26^DTA/DTA^: 69.3%; **Fig. 4B**). Male and female F2 ΔtTreg mice showed no significant differences in mortality (data not shown). Continued backcrossing further exacerbated morbidity, such that none of the F3 ΔtTreg mice lived beyond 14 weeks (**Fig. 7A**).

**Figure 7:**
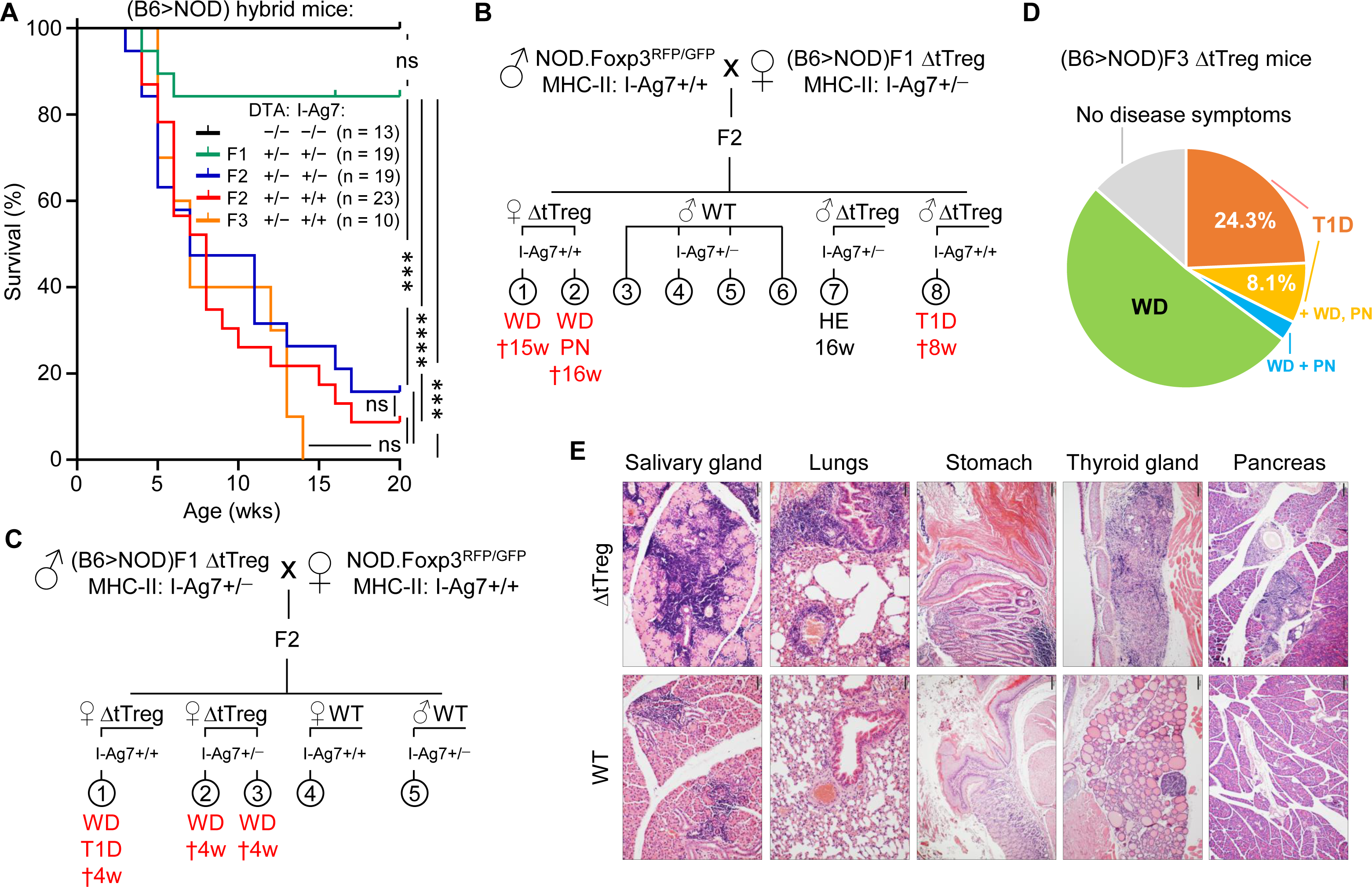
Spontaneous mortality and autoimmune pathology in ΔtTreg mice on a mixed (B6>NOD) background. B6.ΔtTreg mice were backcrossed to NOD.Foxp3^RFP/GFP^ mice for up to 3 generations, as indicated (F1, F2, F3). (B6>NOD) hybrid offspring were analyzed by flow cytometry for the haplotype-specific expression of MHC class II molecules (NOD: I-A^G7^; B6: I-A^b^). **(A)** Kaplan-Meier survival analysis. Cohorts of DTA^+/−^ ΔtTreg mice on a mixed (B6>NOD) background and with heterozygous I-Ag7^+/−^ (F1, n = 19; F2, n = 19) or homozygous I-Ag7^+/+^ (F2, n = 23; F3, n = 10) expression were monitored for the occurrence of spontaneous cases of mortality and morbidity from birth onwards for up to 20 weeks, as indicated. Note that tTreg cell-proficient B6.Foxp3^RFP/GFP^ mice (DTA^−/−^, I-Ag7^−/−^; n = 13) were included for comparison. Log-rank test and Bonferroni correction: ns, not significant; ***p ≤ 0.001, ****p ≤ 0.0001. **(B,C)** Representative pedigrees of **(B)** (NOD>F1) and **(C)** (F1>NOD) ♂x♀matings and health status of resultant F2 offspring. Parental (B6>NOD)F1 mice in **(B,C)** were obtained from independent (B6>NOD) backcross breedings. WD, wasting disease; T1D, type 1 diabetes; PN, peripheral neuropathy; HE, healthy (no disease symptoms); †: age of death in weeks. **(D)** Morbidity of (B6>NOD)F3 ΔtTreg mice (37 males and females from 7 litters). WD: 51.4%, T1D: 24.3%, T1D + WD + PN: 8.1%, WD + PN: 2.7%, no disease symptoms: 13.5%. **(D)** Histological analysis. Organs of 11-13-week-old (B6>NOD)F3 males (I-Ag7^+/+^) were fixed in 4% PFA and cut sections were stained with hematoxylin and eosin. Representative histology showing pronounced leukocytic infiltrations and severe histopathological changes (score: +++) in the salivary gland, lungs, thyroid gland, stomach, and pancreas of a hyperglycemic R26^DTA^ NOD.ΔtTreg F3 mouse (top panels). R26^wt/wt^ NOD.Foxp3^RFP/GFP^ F3 mice (WT, bottom panels) show comparably severe immune infiltrates in the salivary glands, but only marginal (lungs, pancreas) or no (stomach, thyroid gland) autoimmune infiltration in other organs.

In order to account for possible variability of disease pathology on a mixed (B6>NOD) background, we produced independent cohorts of F2 and F3 ΔtTreg mice originating from unrelated (B6>NOD) backcross breedings and parental (B6>NOD)F1 mice. When we monitored the ΔtTreg offspring for *scurfy*-like symptoms, we found that the high incidence of spontaneous mortality depicted in **Fig. 7A** was consistently accompanied by signs of distinct, partially overlapping autoimmune diseases in independent cohorts of F2 ΔtTreg mice (**Fig. 7B,C**). Most prominently, we observed signs of wasting disease (WD; reduced body weight and size, failure to thrive), autoimmune diabetes (hyperglycemia), and peripheral neuropathy (PN; hindlimb paralysis). In the (B6>NOD)F3 generation, ≥ 90% of ΔtTreg mice suffered from either WD (62.2%), T1D (32.4%), and/or PN (10.8%) (**Fig. 7D**). Histopathological analyses indicated massive immune infiltrates of the salivary glands, the lung, the stomach, the thyroid glands, and the pancreas associated with severe tissue damage predominantly affecting thyroid glands and pancreas of NOD.ΔtTreg F3 mice, which correlated with their hyperglycemic state (**Fig. 7E**; top panel). In these experiments, other organs commonly targeted by severe autoimmune responses in Foxp3-deficient mice showed no or only minimal immune infiltrates in NOD.ΔtTreg F3 mice, such as the small intestine or the liver (data not shown). Notably, severe salivary gland autoimmunity could be observed in both ΔtTreg and Foxp3^RFP/GFP^ mice (**Fig. 7E**; left panels), and thus driven by the increased genetic autoimmune risk of the (B6>NOD)F3 background, rather than selective tTreg cell paucity. Other hallmarks of the fatal autoimmune syndrome affecting Foxp3-deficient mice could not be observed, such as skin lesions or scaliness and crusting of the eyelids, ears, and tail (data not shown). Throughout the present study, the manifestation of WD, T1D, and/or PN in tTreg cell-proficient littermates of (B6>NOD) hybrid ΔtTreg mice has also not been observed (**Fig. 7B-E**; **Fig. 8A**; and data not shown).

**Figure 8:**
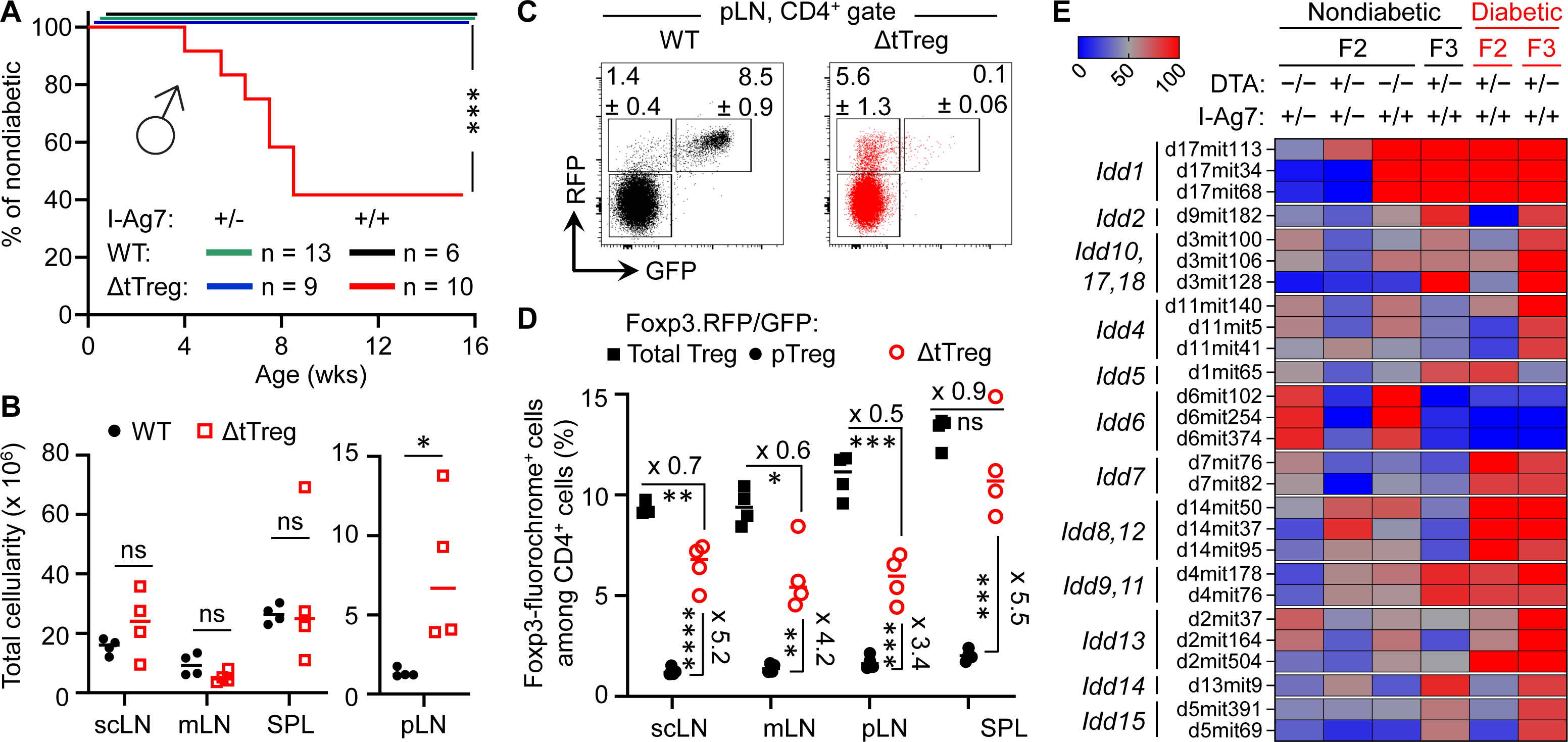
Pancreatic β cell autoimmunity in hybrid (B6>NOD) ΔtTreg mice. Blood glucose levels of 3-week-old F2 ΔtTreg males with heterozygous (I-Ag7^+/−^, n = 9) or homozygous (I-Ag7^+/+^, n = 10) expression of I-Ag7 were monitored once a week. I-Ag7^+/−^ (n = 13) and I-Ag7^+/+^ (n = 6) F2 Foxp3^RFP/GFP^ mice (WT) mice were included for comparison. Note that initially normoglycemic mouse cohorts were selected based on the absence of WD, PN, or other scurfy-like symptoms. Log-rank test: ns, not significant; ***p ≤ 0.001. **(B-D)** Flow cytometric immunophenotyping of (B6>NOD)F2 mice depicted in Fig. 8A. **(B)** Total cellularity of pLN (right) and other peripheral lymphoid tissues (scLN, mLN, spleen) of Foxp3^RFP/GFP^ (closed black circles) and ΔtTreg (open red squares) mice (all 8-week-old I-Ag7^+/+^ males). See **Supplementary Fig. S4A,B** for corresponding numbers of CD8^+^ and CD4^+^ T cells. **(C)** Representative dot plots of Foxp3-driven RFP and GFP expression among CD4-gated cells from pLN from Foxp3^RFP/GFP^ and ΔtTreg mice, and **(D)** cumulative percentages of total Treg (closed black squares) and pTreg (closed black circles) cells of Foxp3^RFP/GFP^ mice, as well as pTreg cells of ΔtTreg mice (open red circles) from peripheral lymphoid tissues, as indicated (all I-Ag7^+/+^). Symbols and horizontal lines in **(B,D)** indicate mean values ± SD of 4 mice per group, as depicted in Fig. 8A. Unpaired t-test: ns, not significant; *p ≤ 0.05, **p ≤ 0.01, ***p ≤ 0.001, **** p ≤ 0.0001. **(E)** Genomic DNA-based *Idd* gene locus analysis in cohorts of (B6>NOD) hybrid mice. Cohorts of F2 Foxp3^RFP/GFP^ mice (WT; I-Ag7^+/−^: n = 17; I-Ag7^+/+^, n = 11); F2 ΔtTreg mice (I-Ag7^+/−^, n = 7; I-Ag7^+/+^, n = 5), and F3 ΔtTreg mice (nondiabetic I-Ag7^+/+^, n = 8; diabetic I-Ag7^+/+^, n = 5) were subjected to genomic PCR for *Idd* gene analysis. The heatmap shows the distribution of a selected set of *Idd* loci among different experimental groups, as indicated. For this, the percentage of mice homozygous for the respective NOD *Idd* gene locus was calculated and expressed as color code. An overview of the complete data set is provided in **Supplementary Fig. S5**. Note that the R26-DTA transgene of ΔtTreg mice is embedded within the *Idd6* gene locus, resulting in a marked underrepresentation of *Idd6* in F2 and F3 ΔtTreg mice.

### Pancreatic β cell autoimmunity in(B6>NOD) hybrid ΔtTreg mice

While Foxp3^+^ Treg cell-deficient NOD mice fail to develop insulitis and overt diabetes (Chen et al., 2005), the data depicted in **Fig. 7** provided the first indications that selective tTreg cell paucity can promote severe insulitis and overt diabetes (32.4%; **Fig. 7B-E**) in both males and females, despite incomplete backcrossing onto the NOD background. However, more definite conclusions on the role of tTreg cells in controlling pancreatic β cell autoimmunity were hampered by the overall early onset of high morbidity and mortality (**Fig. 7A**). In fact, three diabetic F3 NOD.ΔtTreg mice additionally exhibited signs of WD and PN (**Fig. 7D**), suggesting that some (B6>NOD) hybrid ΔtTreg mice may succumb to death before the diabetes diagnosis. We therefore tracked blood glucose levels in cohorts of 3-week-old, initially normoglycemic F2 NOD.ΔtTreg mice that showed no signs of WD, PN, or other *scurfy*-like symptoms (**Fig. 8A**). In our colony of conventional NOD mice, the first diabetes cases become apparent at approximately 12 weeks of age and continuously increase to an incidence of 70-90% in females and 0-20% in males within 30 weeks of age (Petzold et al., 2010; Watts et al., 2021). Whereas in (B6>NOD)F2 ΔtTreg mice, selective tTreg cell paucity unleashed a particularly severe form of T1D: > 50% of males rapidly progressed to overt diabetes within < 8 weeks after birth (**Fig. 8A**), despite the usually observed female sex bias and kinetics difference in the NOD model (Yurkovetskiy et al., 2013). Flow cytometry-based MHC class II haplotyping indicated that diabetes manifestation in F2 ΔtTreg mice correlated with homozygous expression of the diabetogenic MHC class II molecule I-Ag7 of the NOD genetic background (*Idd1*), whereas mice co-expressing I-Ab of B6 origin remained normoglycemic during the observation period (**Fig. 8A**). In contrast, high mortality (**Fig. 7A**) and the manifestation of WD (**Fig. 7C**) was independent of homozygous I-Ag7 expression.

In line with the absence of severe systemic autoimmune responses, pLNs of F2 ΔtTreg mice showed clear signs of lymphadenopathy, whereas the size of non-draining LNs (scLNs, mLNs) and spleen did not significantly differ between tTreg cell-deficient and -proficient (B6>NOD)F2 mice (**Fig. 8B**). Consistently, numbers of CD8^+^ (**Supplementary Fig. S4A**) and CD4^+^ (**Supplementary Fig. S4B**) T cells were selectively increased in pLNs of F2 ΔtTreg mice.

Consistent with our data in B6.ΔtTreg mice (**Fig. 1D,E**), efficient intrathymic tTreg cell ablation in (B6>NOD)F2 ΔtTreg mice (**Supplementary Fig. S4C**) was accompanied by a significant, up to 5.5-fold increase in the percentage of RFP^+^GFP^−^ pTreg cells among CD4-gated T cells in peripheral lymphoid tissues (**Fig. 8C,D**). However, in contrast to B6.ΔtTreg mice (**Supplementary Fig. S1A**), the population size of pTreg cells in (B6>NOD)F2 ΔtTreg mice only partially compensated for the numerical impairment of the overall Treg cell pool in the absence of tTreg cells (**Fig. 8D**). Additionally, thymic cellularity (**Supplementary Fig. S4D**) and numbers of T cell developmental stages (**Supplementary Fig. S4E**) were consistently reduced in (B6>NOD)F2 ΔtTreg mice, as compared to in (B6>NOD)F2 Foxp3^RFP/GFP^ mice, probably due to increased hyperglycemia-induced stress and/or systemically elevated inflammatory cytokine levels.

### Contribution of NOD *Idd* loci to diabetes in(B6>NOD) hybrid ΔtTreg mice

We found that the manifestation of overt diabetes was restricted to I-Ag7^+/+^ (B6>NOD) hybrid ΔtTreg mice (**Fig. 7B-E**; **Fig. 8A**), consistent with the requirement of *Idd1* homozygosity for high penetrance of diabetes susceptibility in the NOD model (Hattori et al., 1986). In fact, *Idd1* was shown to confer most of the diabetes risk (Ikegami et al., 2004), but not to be sufficient to precipitate diabetes in Foxp3^+^ Treg cell-proficient NOD mice (Ikegami et al., 2004). We hypothesized that the early manifestation of diabetes in I-Ag7^+/+^ F2 ΔtTreg males with high penetrance (**Fig. 8A**) was driven by the acquisition of one or more additional, non-MHC-linked *Idd* loci. As expected, after only two backcross generations, PCR-based genomic *Idd* gene analysis (**Supplementary Fig. S5**) indicated that the majority of *Idd* genes included in our survey was dispersible for diabetes development in I-Ag7^+/+^ F2 ΔtTreg mice (*Idd2*, *Idd3/Idd10/Idd17/Idd18*, *Idd4, Idd14, Idd15*) (**Supplementary Fig. S5A**). This included *Il2* gene polymorphisms (encoded by *Idd3*), which play an important role in the reduced IL-2 receptor signaling strength received by Treg cells in conventional, tTreg cell-proficient NOD mice, resulting in their functional deficiency (Tang et al., 2008; Long et al., 2011). Additionally, *Idd6* was markedly underrepresented in (B6>NOD)F2 (**Supplementary Fig. S5A; Fig. 8E**) and F3 **(Supplementary Fig. S5B; Fig. 8E)** ΔtTreg mice, as compared to their Foxp3^RFP/GFP^ littermates, which can be attributed to the genomic localization of the R26-DTA transgene of ΔtTreg mice within the *Idd6* gene locus (http://www.informatics.jax.org/). Other *Idd* gene loci initially underrepresented in diabetic F2 ΔtTreg mice were found to be enriched after continued backcrossing (*Idd2, Idd4, Idd3/Idd10/Idd17/Idd18, Idd13.1/.2, Idd14, Idd15)* (**Supplementary Fig. S5B; Fig. 8E**).

In addition to *Idd1*, a set of 5 *Idd* gene loci (*Idd5.1*, *Idd7*, *Idd8/Idd12, Idd9.1/.2, Idd13.3*) was detectable in ≥ 80% of diabetic F2 ΔtTreg mice (**Fig. 8E**), but was not sufficient to promote diabetes in tTreg cell-proficient I-Ag7^+/+^ (B6>NOD)F2 littermates (**Fig. 8A**; and data not shown). Interestingly, this rather small set of *Idd* loci was primarily characterized by harboring genes with well-known functions in the development, survival/maintenance, function of pancreatic β cells [*Idd7*, *Idd8*, *Idd12* (Gao et al., 2008)(Itoh et al., 2003; Goulley et al., 2007; Hussain, 2007; Schumann et al., 2007)] and Foxp3^+^ Treg cells (*Idd5*, *Idd7, Idd9.1/.2, Idd13.3*). In particular, several annotated genes located in *Idd5* [*Cd28*, *Icos*, *Ctla4* (Greve et al., 2004; Wicker et al., 2004); *Pdcd1* (Simpfendorfer et al., 2015); *Irs1*, *Stat1* (Ligons et al., 2009); *Ikfz2* (Hunter et al., 2007)] and the *Idd9* gene locus [(*Cd30, Tnfr2, Cd137* (Lyons et al., 2000; Forsberg et al., 2019)*; p110δ, mTOR* (Hamilton-Williams et al., 2013; Ferreira et al., 2014)] play key roles in various facets of Foxp3^+^ Treg cell biology (Sharpe and Freeman, 2002; Bour-Jordan and Bluestone, 2009; Chapman and Chi, 2014; Chougnet and Hildeman, 2016; Gianchecchi and Fierabracci, 2018), many of which belong to the shared transcriptional signatures of tissue-type Treg (Delacher et al., 2017, 2020; Miragaia et al., 2019) and pTreg (Petzold et al., 2014) cells. Furthermore, and directly relevant to the autoimmune mechanisms underlying pancreatic β cell destruction, *Idd7* (containing *Nfkbid*) has been implicated in modulating diabetogenic CD8^+^ T cell deletion in the thymus (Presa et al., 2018) and numbers and suppressor function of Foxp3^+^ Treg cells in peripheral tissues (Dwyer et al., 2022). Lastly, several annotated genes encoded in *Idd13* [*B2m* (Serreze et al., 1998; Hamilton-Williams et al., 2001)*; Mertk* (Wallet et al., 2009)*; Bcl2l11/Bim, Id1, Smox, Pdia3* (Liston et al., 2007)] play key roles in development and activation of diabetogenic T effector cells (Serreze et al., 1998; Hamilton-Williams et al., 2001), including negative thymic selection (Liston et al., 2007; Wallet et al., 2009; Dugas et al., 2014).

Overall, these findings in ΔtTreg mice are consistent with a scenario, in which pTreg cell-mediated maintenance of immunological tolerance to pancreatic β cells can be abrogated by the acquisition of a limited set of *Idd* risk loci, some of which unfold their diabetogenic activity directly in pTreg cells. In support of this interpretation, comparative flow cytometry-based immunophenotyping revealed a correlation of some detected *Idd* loci and differential protein expression in pTreg cells of F2 ΔtTreg mice, as compared to pTreg cells in tTreg cell-proficient Foxp3^RFP/GFP^ littermates (**Fig. 9**). This included the absence of *Idd3* (including *Il2*) and markedly increased expression levels of CD25 on pTreg cells from pLNs of diabetic F2 ΔtTreg mice, as compared to F2 Foxp3^RFP/GFP^ (**Fig. 9A,C**; left panels). Relevant to their high diabetes susceptibility, the acquisition of *Idd5.1* (Greve et al., 2004) by F2 ΔtTreg mice correlated with increased expression levels of ICOS (*Icos*) and PD-1 (*Pdcd1*) on pTreg cells (**Fig. 9A,C**), and the accumulation of an unusual ICOS^+^PD-1^high^ pTreg cell subset in F2 ΔtTreg mice, but not in Foxp3^RFP/GFP^ littermate controls (**Fig. 9D**). The expression levels of other Treg signature proteins remained largely unaffected (*e.g.*, Nrp-1, GITR, KLRG1) (**Fig. 9B**). Importantly, functional incapacitation of PD-1 in gene-targeted mice (Salama et al., 2003; Kroner et al., 2009; Wang et al., 2009) and human patients treated with blocking Abs (Tucker et al., 2021) can result in overt autoimmune responses, including T1D (Ansari et al., 2003; Wang et al., 2005; Keir et al., 2006; Paterson et al., 2011; Tucker et al., 2021), indicating a primarily inhibitory function of PD-1 expression in immune effector cells. However, independent lines of evidence in mice have pointed towards a diabetogenic role of PD-1 expression on Foxp3^+^ Treg cells, which included diabetes amelioration in congenic B10.Idd5^+^ and Idd5.1^+^ NOD mice (Wicker et al., 2004; Lamhamedi-Cherradi et al., 2001), diabetes protection of NOD mice with Foxp3^+^ Treg cell-specific PD-1 deletion (Tan et al., 2021), and an inverse correlation of PD-1 expression with the expression of Foxp3 and Foxp3^+^ Treg cell function (Wong et al., 2013; Tan et al., 2021). Clearly, future studies are warranted to further dissect the differential function of PD-1 on T effector and Foxp3^+^ Treg cells, including pTreg cells.

**Figure 9:**
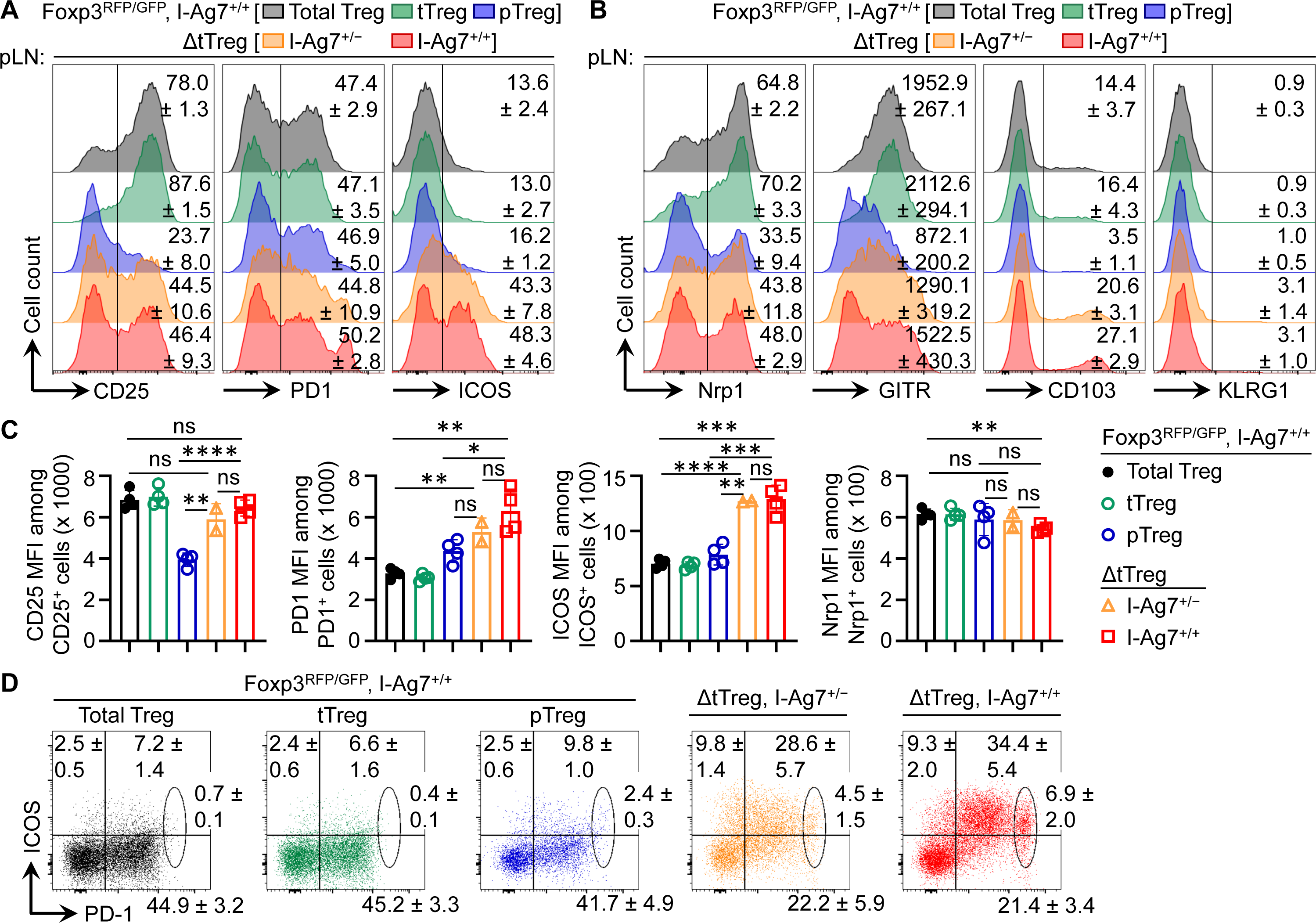
Flow cytometry-based immunophenotyping of pTreg cells in (B6>NOD)F2 ΔtTreg mice. Expression analysis of selected Treg cell signature proteins on gated Treg cell subsets, as indicated. Surface markers with **(A)** upregulated and **(B)** unchanged expression on pTreg cells from F2 ΔtTreg mice, as compared to pTreg cells from F2 Foxp3^RFP/GFP^ mice. Numbers in histograms indicate mean percentages of cells ± SD within the respective gate or quadrant, with the exception of GITR in **(B)** showing mean fluorescence intensity (MFI) of the fluorochrome-conjugated mAb **(C)** Quantification of indicated marker expression based on MFI of fluorochrome-conjugated mAbs. Symbols and bars represent individual mice and mean values ± SD, respectively. **(D)** Expression of ICOS and PD-1 on pTreg cells from pLNs of Foxp3^RFP/GFP^ mice (I-Ag7^+/+^) and ΔtTreg mice (I-Ag7^+/−^ or I-Ag7^+/+^), as indicated. Numbers in dot plots indicate mean percentages of cells ± SD within the respective quadrant or gate. Note that gated populations of total Treg cells and tTreg cells from Foxp3^RFP/GFP^ mice were included for comparison. nt. Data are from a single experiment (n = 4). Unpaired t-test: ns, not significant; *p ≤ 0.05, **p ≤ 0.01, ***p ≤ 0.001, **** p ≤ 0.0001.

## DISCUSSION

As the functional heterogeneity of pTreg and tTreg cells promises to enable the subset-specific therapeutic manipulation of their activity in various clinical settings, it will be important to define their exact roles in establishing and maintaining peripheral immune homeostasis. The selective ablation of the development of pTreg cells (Zheng et al., 2010; Josefowicz et al., 2012; Campbell et al., 2018) and tTreg cells, as done here, represents a considerable improvement over previous experiments relying on Foxp3-deficient mice and their reconstitution by adoptive Treg cell transfer, allowing the *in vivo* consequences to be analyzed under near-physiological conditions, including minimal autoimmune perturbations. In fact, some of the pathology observed in Foxp3-deficient mice has been attributed to the enhanced thymic export and peripheral accumulation of Treg cell-like ‘wanna-be’ CD4^+^ T cells with self-reactive specificities and distinct pathological properties (Kuczma et al., 2009; Wyss et al., 2016), rather than the mere absence of a functional Foxp3^+^ Treg cell pool. Additionally, some defects of Foxp3-deficient mice (*e.g.*, defective lympho-hematopoiesis) are refractory to adoptive Treg cell therapy, even when total CD4^+^ T cell populations were used (Riewaldt et al., 2012). Here we have analyzed how selective tTreg cell paucity, which was achieved by intrathymic tTreg cell ablation while preserving pTreg cell generation, impinges on peripheral immune homeostasis in non-autoimmune and autoimmune-prone mice. Our data in B6.ΔtTreg mice reveal the ability of pTreg cells to establish immune homeostasis after birth, maintain immune tolerance in young mice, and constrain catastrophic autoimmune responses during aging in the majority of B6.ΔtTreg mice. Consistently, neonatal transfer of total CD4^+^ T cell populations, which had been depleted of tTreg cells, ameliorated clinical signs of Foxp3 deficiency in *scurfy* recipient mice. The manifestation of some mild disease symptoms (moderate growth delay and mild exfoliative dermatitis) can probably be attributed to an initial lag phase after tTreg cell-depleted CD4^+^ T cell transfer, which is required for seeding and proliferative expansion of pre-formed pTreg cells, and the lymphopenia-driven *de novo* generation of Foxp3^+^ pTreg cells (Haribhai et al., 2011).

Besides the absence of *scurfy*-like symptoms, several additional observations in the B6.ΔtTreg model further support our interpretation that physiologic pTreg cell populations can efficiently constrain autoimmune responses in the absence of tTreg cells. This includes an overall normal size of peripheral lymphoid tissues and T effector cell compartments (numbers, activation state, inflammatory cytokine production, *etc.*), as well as unperturbed lympho-hematopoiesis, representing a particularly sensitive indicator for the absence of ongoing (auto)immune responses. Many organs of B6.ΔtTreg mice, which are commonly targeted by severe autoimmune destruction in Foxp3-deficient mice, show no or only mild immune infiltrations not accompanied by any appreciable tissue destruction. Thus, the underlying cause promoting the occurrence of spontaneous deaths from an age of > 7 weeks onwards has remained less clear but may involve the exacerbation of chronic, low-level inflammation in individual organs, such as the thyroid gland or the lungs (**Fig. 4A**), rather than multi-organ autoimmunity observed in Foxp3-deficient models of complete Treg cell deficiency. Interestingly, extending our histopathological analyses of the lungs (**Fig. 4A**) to the upper respiratory tract of ΔtTreg mice that presented with reduced body weight provided first evidence for unexpected, severe inflammatory changes in the area of the nasal and oral cavities, pointing towards decreased food intake as a possible reason underlying a reduced body weight and morbidity of this particular disease subphenotype (data not shown).

Considering the age-dependent increase in spontaneous deaths (**Fig. 4B**), the abrogation of immune homeostasis in individual ΔtTreg mice is likely to involve immunological and/or environmental cues (Sefik et al., 2015; Russler-Germain et al., 2017; Campbell et al., 2018), which are subject to age-related changes. This may include differences in the exposure to antigens derived from the diet and commensal microbiota promoting the physiologic induction of pTreg cells (Lathrop et al., 2011; Arpaia et al., 2013; Sefik et al., 2015; Kim et al., 2016). Our efforts to further analyze the immune events associated with the age-related impairment of peripheral immune homeostasis in ΔtTreg mice have been hampered by the relatively low incidence of mortality, in conjunction with rapid disease progression. In-depth flow cytometry-based immunophenotyping failed to reproducibly reveal age-related changes in the peripheral immune effector compartments of ΔtTreg mice, including CD4^+^ and CD8^+^ T effector compartments (**Fig. 5C-E**). While this could be taken as an indication for quantitative and/or qualitative changes affecting the pTreg cell compartment, our analyses have not provided any evidence for an age-related reduction of pTreg cell numbers or phenotypic changes in the peripheral pTreg pool. In contrast, we found that the increased pTreg cell population size largely compensated for the numerical impairment of the overall Treg cell pool in adult ΔtTreg mice (**Fig. 1D,E; Supplementary Fig. S1**), which also holds true for ΔtTreg mice that were affected by reduced body weight and thymic atrophy (**Fig. 2B**).

At present, we can only speculate on whether the pTreg cell niche in peripheral lymphoid tissues of ΔtTreg mice is replenished early in life and then maintained by proliferative expansion of pre-formed Treg cells, or whether continuous incorporation of newly formed cells is required to maintain a peripheral pTreg cell pool and immune homeostasis. This raises the possibility that the observed age-dependent increase in spontaneous mortality and morbidity is, at least in part, associated with a reduced efficiency in pTreg cell generation. In fact, pTreg cells are thought to be mainly, if not exclusively drawn from initially naïve CD4^+^ T cells (Schallenberg et al., 2013). However, rates of thymic export of newly formed CD4^+^ T cells to peripheral sites of pTreg cell generation continuously decrease during aging and involution of the thymus (Hale et al., 2006; Fink and Hendricks, 2011), but also during thymic atrophy due to chronic inflammatory stress (**Fig. 2B, Supplementary Fig. S4D**). Consistently, immediate CD25^high^Foxp3^−^ pTreg cell precursors residing in peripheral lymphoid tissues of nonmanipulated mice are strongly enriched among recent thymic emigrants (Schallenberg et al., 2010).

Overall, our findings are consistent with a scenario, in which pTreg cells in peripheral lymphoid tissues of B6.ΔtTreg mice acquire a highly activated phenotype and increased suppressor function to cope with latent, chronic autoimmune responses due to the absence of tTreg cells. This intricate equilibrium can get out of control even by subtle age-related immunological and/or microenvironmental changes, which then tip the balance in favor of fatal autoimmunity. This may include changes in the commensal microbiota and qualitative differences among the pTreg cell pool, *e.g.*, reduced rates of pTreg cell *de novo* generation, in conjunction with proliferative pTreg cell expansion narrowing the TCR repertoire.

This interpretation was further corroborated by the dramatically increased mortality associated with the early onset of severe autoimmune diseases that could be observed in ΔtTreg mice after only two backcross generations onto the autoimmune-prone NOD background. Here we focused our analysis on pancreatic β cell autoimmunity, as T1D is considered a paradigmatic autoimmune disease for the application of Treg cell-based therapies to prevent or interfere with ongoing autoimmune destruction, although the main regulator(s) of pancreatic β cell autoimmunity hasn’t been identified yet. Lastly, Foxp3-deficient NOD mice with a polyclonal CD4^+^ T cell repertoire fail to present with insulitis and overt diabetes before they succumb at 3 weeks to severe inflammatory infiltration in multiple organs (Chen et al., 2005), precluding NOD.Foxp3-deficient mice as an experimental model to study the role of Treg cells in the autoimmune b cell protection. Our data show that the acquisition of a small set of *Idd* risk loci, many of which encode genes with well-known functions in Treg cell biology, is sufficient to precipitate a particularly severe form of autoimmune diabetes in ΔtTreg mice on a mixed (B6>NOD) background. In this context, it is of interest to note that our complementary studies in ΔpTreg mice on a (B6>NOD)F5 background haven’t provided evidence for a protective role of pTreg cells in the control of β cell autoimmunity (D.M.Z. and K.K., unpublished observation). Our observations in (B6>NOD) hybrid mice with selective tTreg cell paucity, in conjunction with previous experiments in Foxp3.CNS1^−/−^ with impaired pTreg cell development (Holohan et al., 2019) indicate that tTreg cells are key regulators of β cell autoimmunity in the NOD model. Clearly, future experiments are warranted using Treg cell-subset-deficient mice on a pure NOD background to provide a more definite answer on the role of tTreg and pTreg cells in the control of β cell autoimmunity.

In conclusion, ΔtTreg and ΔpTreg mice offer to directly analyze the individual roles of tTreg and pTreg cells, respectively, in the control of immune homeostasis and organ-specific autoimmunity under near-physiologic conditions, which will facilitate future studies on the functional heterogeneity of the mature Treg cell pool. Besides autoimmune diseases, of particular interest will be to dissect their subset-specific contributions to non-immune functions that have recently been attributed to tissue-type Treg cells, which include facilitating homeostasis and regeneration of nonlymphoid tissues.

## ACKNOWLEDGMENTS

The authors would like to thank Sonja Schallenberg (CRTD) for helpful discussions and constructive advice; Lena Biedermann (CRTD) for technical assistance; and Katja Bernhardt and Anne Gompf (CMCB Flow Cytometry Core Facility) for expert help in flow cytometry and cell sorting. Additionally, the authors are greatly indebted to Abdel Gdoura and Peter van Endert (Université Paris Cité, INSERM) for sharing their longstanding expertise in *Idd* gene analysis. We also would like to thank Basak Torun for critically reading of our manuscript

## AUTHOR CONTRIBUTIONS

Acelya Yilmazer designed, performed and analyzed the experiments, and contributed to the data interpretation and writing of the manuscript. Dimitra Maria Zevla, Rikke Malmkvist, Carlos Alejandro Bello Rodríguez, Pablo Undurraga, Emre Kirgin, Olivia Kershaw, and Marie Boernert performed experiments. David Voehringer and Susan Schlenner contributed to the research design and the analysis and interpretation of data. Additionally, Susan Schlenner developed the conditional Foxp3-STOP allele and contributed to the writing of the manuscript. Karsten Kretschmer conceived the research, guided its design, analysis, and interpretation, and wrote the manuscript.

---

**Supplementary Figure S1: Treg cell frequencies in peripheral lymphoid tissues and blood.** Percentages of CD4-gated Foxp3-fluorochrome^+^ Treg cells in peripheral lymphoid tissues (scLNs, mLNs, and SPL) and peripheral blood of **(A,B)** 13-22-week-old (13 mice per group) and **(C)** 52-56-week-old (3-4 mice per group) male mice, as indicated. (Foxp3^RFP/GFP^ mice: RFP^+^GFP^⎻^ pTreg, filled black circles; RFP^+^GFP^+^ tTreg: filled black squares; total RFP^+^ Treg: open black diamonds; ΔtTreg mice: RFP^+^GFP^⎻^ pTreg, open red circles; ΔpTreg mice: GFP^+^ tTreg, open green squares). Symbols and horizontal lines represent individual mice and mean values, respectively. Unpaired t-test: ns, not significant; *p ≤ 0.05, **p ≤ 0.01, ***p ≤ 0.001, ****p ≤ 0.0001.

**Supplementary Figure S2: Normal B lymphopoiesis in adult ΔtTreg and ΔpTreg mice.** Flow cytometry of B cell development in bone marrow (BM) and SPL of ΔtTreg (n = 7), ΔpTreg (n = 6), and Foxp3^RFP/GFP^ (n = 7) mice at 10-15 weeks of age. **(A)** Total cellularity of BM (top) and SPL (bottom). **(B-E)** B cell development. **(B)** Representative flow cytometry and **(C)** numbers of B220^+^c-kit^+^ Pro/Pre-B-I cells (top), as well as immature B220^low^IgM^+^ and mature/recirculating B220^high^IgM^+^ B cells (bottom) in BM. **(D)** Representative flow cytometry and **(E)** numbers of immature IgD^low^IgM^high^ (left) and mature IgD^high^IgM^low^ (right) B cells in SPL. Numbers in dot plots in **(B,D)** represent mean percentages of cells ± SD within the respective gate. Symbols and bars in **(A,C,E)** represent individual mice (WT: black circles; ΔtTreg: open red squares; ΔpTreg: open green triangles) and mean values, respectively. Unpaired t-test: ns, not significant. Data are from two independent experiments with 6-7 mice per group.

**Supplementary Figure S3: pTreg cells recirculate to the thymus of ΔtTreg mice.** Tracking thymic recirculation of mature Treg cells to the thymus. **(A)** Representative dot plots of CD62L and Foxp3-driven RFP reporter expression among CD4SP-gated cells in the thymus of Foxp3^RFP/GFP^ (WT, n = 5) and ΔtTreg (n = 3) males at 17-22 weeks of age. **(B)** Histograms show the expression of GFP and surface markers on gated cell populations, as indicated (blue: RFP^int^CD62L^high^; yellow: RFP^high^CD62L^high^; red: RFP^high^CD62L^⎻^). Note that the RFP^high^CD62L^high^ CD4SP compartment is readily detectable in Foxp3^RFP/GFP^ mice (top panels) but below the detection limit in ΔtTreg mice (bottom panels) due to selective DTA-mediated deletion. Numbers in dot plots in **(A)** and histograms of GFP and CD25 expression in **(B)** represent mean percentages of cells ± SD within the respective gate. Numbers in histograms of CD69 and CD44 expression represent mean fluorescence intensities (MFI) of marker expression ± SD. Data are from a single experiment representative of 3 experiments performed (3-6 mice per experiment).

**Supplementary Figure S4: Peripheral T cell numbers and intrathymic tTreg cell ablation in (B6>NOD)F2 ΔtTreg mice.** Numbers of **(A)** CD8^+^ T cells and **(B)** CD4^+^ T cells from pLN (right) and other peripheral lymphoid tissues (scLN, mLN, spleen) of Foxp3^RFP/GFP^ (WT, closed black circles) and ΔtTreg (open red squares) mice (all 8-week-old I-Ag7^+/+^ males), corresponding to total organ cellularity depicted in **Fig. 8B. (C-E)** Intrathymic tTreg cell ablation and thymopoiesis. **(C)** Representative flow cytometry of CD4 and CD8 expression (left) and CD25 and Foxp3-driven GFP expression (right) among FSC/SSC-gated cells from the thymus (THY) of 8-week-old Foxp3^RFP/GFP^ and ΔtTreg male mice. Numbers in dot plots indicate mean percentages of cells ± SD within the respective gate. **(D)** Total thymic cellularity, as well as **(E)** numbers of DP, CD4SP, and CD8SP thymocytes, as indicated. Symbols and bars represent individual mice and mean values ± SD, respectively. Data in **(C-E)** are from a single experiment (n = 4), corresponding to the data depicted in **Fig. 8B-D**. Unpaired t-test: ns, not significant; *p ≤ 0.05, **p ≤ 0.01.

**Supplementary Figure S5: Genomic DNA-based *Idd* gene locus analysis of individual (B6>NOD) hybrid mice.** Cohorts **(A)** of F2 Foxp3^RFP/GFP^ mice (nondiabetic WT; I-Ag7^+/−^: n = 17; nondiabetic I-Ag7^+/+^, n = 11); F2 ΔtTreg mice (nondiabetic I-Ag7^+/−^, n = 7; nondiabetic, I-Ag7^+/+^, n = 4; diabetic I-Ag7^+/+^, n = 5), and **(B)** of F3 Foxp3^RFP/GFP^ mice (nondiabetic WT; I-Ag7^+/+^, n = 5) and F3 ΔtTreg mice (nondiabetic I-Ag7^+/+^, n = 8; diabetic I-Ag7^+/+^, n = 5) were subjected to genomic PCR for *Idd* gene analysis. The NOD *Idd* status (blue: absent; grey: heterozygous; red: homozygous) of all 36 loci (including subloci) is shown in individual male (♂) and female (♀) mice for each experimental group, as indicated. Note that the R26-DTA transgene of ΔtTreg mice is embedded within the *Idd6* gene locus, resulting in a marked underrepresentation of *Idd6* in F2 and F3 ΔtTreg mice.

## Supplementary Materials and Methods

The following primers were used for *Idd* loci analyses:

d17mit113: 5’-TCT GTC TCC TCC GTA CTG GG, 5’-GTC AAT AAG TTC AAT CAC TGA ACA CA;

d17mit34: 5’-TGT TGG AGC TGA ATA CAC GC, 5’-GGT CCT TGT TTA TTC CCA GTA CC;

d17mit68: 5’-GTC CTG ACA TCA TGC TTT GTG, 5’-CTA CCG TTT GGA AGG CTG AG;

d9mit91: 5’-TTC TGG GGC AGA GAC CAG, 5’AGA AGG TGG CAG GGG TAG TT;

d9mit208: 5’-GCC TCT CTT TCT TTA AAC ACT TTA AG, 5’-CCT CCA CAC ACC TGT TTG TG;

d9mit182: 5’-GTG AAA TTG GTT ATG TAA ATG TCT GA, 5’-GAG ATG ACT AGG GTG AAC TGG G;

d3nds6: 5’-GTG GGA GTG TGT GCA AAA GAC, 5’-CAG AAT AGG TGA TTA GGT GGT TAT;

d3mit100: 5’-CCT GAT GAC TCT GCG TGT GT, 5’-ATA CCA GTG TTC TCC CCA ACC;

d3mit106: 5’-ACT TGT GCA TGG TGT GTA TGC, 5’-TGT GAT GGC ACC TTT GGT AA;

d3mit128: 5’-AAT AAA GGA AGA TGT CAT CTC AGT ACA, 5’-GAT GGG ATG GGA TGG GAT;

d11mit140: 5’-GCA TTT ACT TGA TTG ATT GTT TGC, 5’-ACC CAA TGC CTG CCT CTA C;

d11mit5: 5’-TTC TGT GAG CCT GGA GGA GT, 5’TAC AGG ACT AGT TTC CAT TTG GG;

d11mit41: 5’-CTG CTA AAG TGG GGT TAA ATG C, 5’-CGA CTG AGC AAG TTG TAT TTC TG;

d1mit65: 5’-CTA ACC CCT ATA CAC ATA CTG CCC, 5’-CCG TTC AGA CTT GAA TAC AGA CC;

d1mit18: 5’-TCT GGT TCC AGG CTT GAT TC, 5’-TCA CAA GTG AGG CTC CAG G;

d1mit8: 5’-CTG AAA ATC GTC CCT TGA CC, 5’-CAG GAG CAT GAA ATG GGG AT;

d1mit30: 5’-TGA ACC ATC ACC ATG CTG TT, 5’-TGG GCT GCG TTT CTA AGG;

d6mit102: 5’-CCA TGT GGA TAT CTT CCC TTG, 5’-GTA TAC CCA GTT GTA AAT CTT GTG TG;

d6mit254: 5’-AGT GTC CCT AGG GGG TGG, 5’-GGG GCC TTA GAG GTA GCA AC;

d6mit374: 5’-TTC TGG CTC TTA ACA GTC TGT CC, 5’-TAC ATA TGC CAA TGA TAT TCT CCC;

d7mit76: 5’-CAT GAG CAC GTG GAG AAA GA, 5’-CGT GGA AAC CTG ATA AAC TGA;

d7mit82: 5’-GGA CAC GGT GTC CAT CAA G, 5’-CTG AGT AGA AAG CAT GTG GGG;

d14mit50: 5-’GAG GGG GAA TCC TAG TGC TC, 5’-AGC AAA GCC CTA TCC ACA TG;

d14mit37: 5’-GTC GAT GGA TGA CTG CTG C, 5’-CAT GGG GAC TCA GGA GAT TG;

d14mit95: 5’-TAT TTT TAA GTC AGT ATA CAC ATG CGC, 5’-TTA TCC AAG TGT ATT TAA AGA AGA GGC;

d4mit178: 5’-GCC CTG AAG GTA AAT CAG TAA CT, 5’-GCT CAG GAG GTA CAT TGC CT;

d4mit76: 5’-TGA AGG AAC CTG AAG CAA GG, 5’-ACC TCC CAG GAG TGT CCA G;

d4mit233: 5’-TGG TCA TGT GTG TCC ATG C, 5’-ACT TCA TGT AGC CAG GTG GG;

d2mit37: 5’-TGT GCA AGC CAG AAA AGT TG, 5’-GAA GGG GAT TGT AAA TTG GTA CC;

d2mit164: 5’-TCT CTG CTA ATT AAG TTG AAG AGT GC, 5’-ACC AGT GTG TGT TTG TAT GAT GTG;

d2mit504: 5’-ATT TCA CAA GTC TTC CCC CC, 5’-TGA AAC ACA AAT GAG CAA CTA CG;

d13mit16: 5’-CCA GCT GAA GGC TTA CTC GT, 5’-AAA GTT AGA ATC AGC CAT TCA AGG;

d13mit61: 5’-TGC TCC AAT ACA ACA AGG TCC, 5’-CCA GCC AAG GTG TGT TGA C;

d13mit9: 5’-GGG TTC CAG ATT GAG TGG AA, 5’-TTG CCA AAG TGT CAA AAT CA;

d5mit391: 5’-AAT AAG AAA ATT CCA CCA AGT CTA CA, 5’-CTT GAT GGG TCT GAT GCC TT;

d5mit69: 5’-CCA GCC TTT CTG GAG TGA AG, 5’-ACC ATG GCA GAA AGC AGT TT.

