## Supplementary figures for "Selective ablation of thymic and peripheral Foxp3^+^ regulatory T cell development"

**Supplementary Figure S1: Treg cell frequencies in peripheral lymphoid tissues and blood.** Yilmazer *et al.*

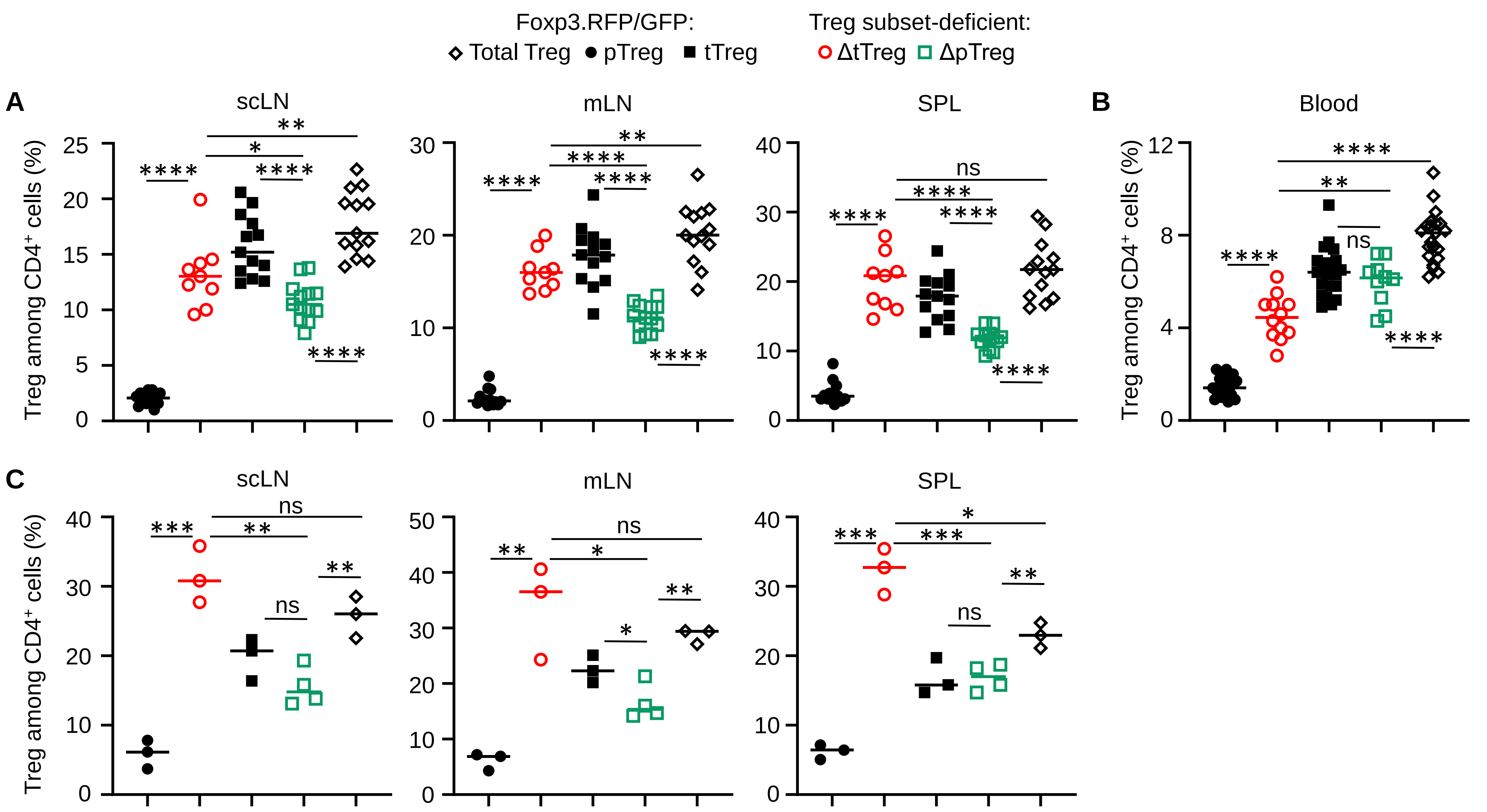

**Supplementary Figure S2: Normal B lymphopoiesis in adult ΔtTreg and ΔpTreg mice.** Yilmazer *et al.*

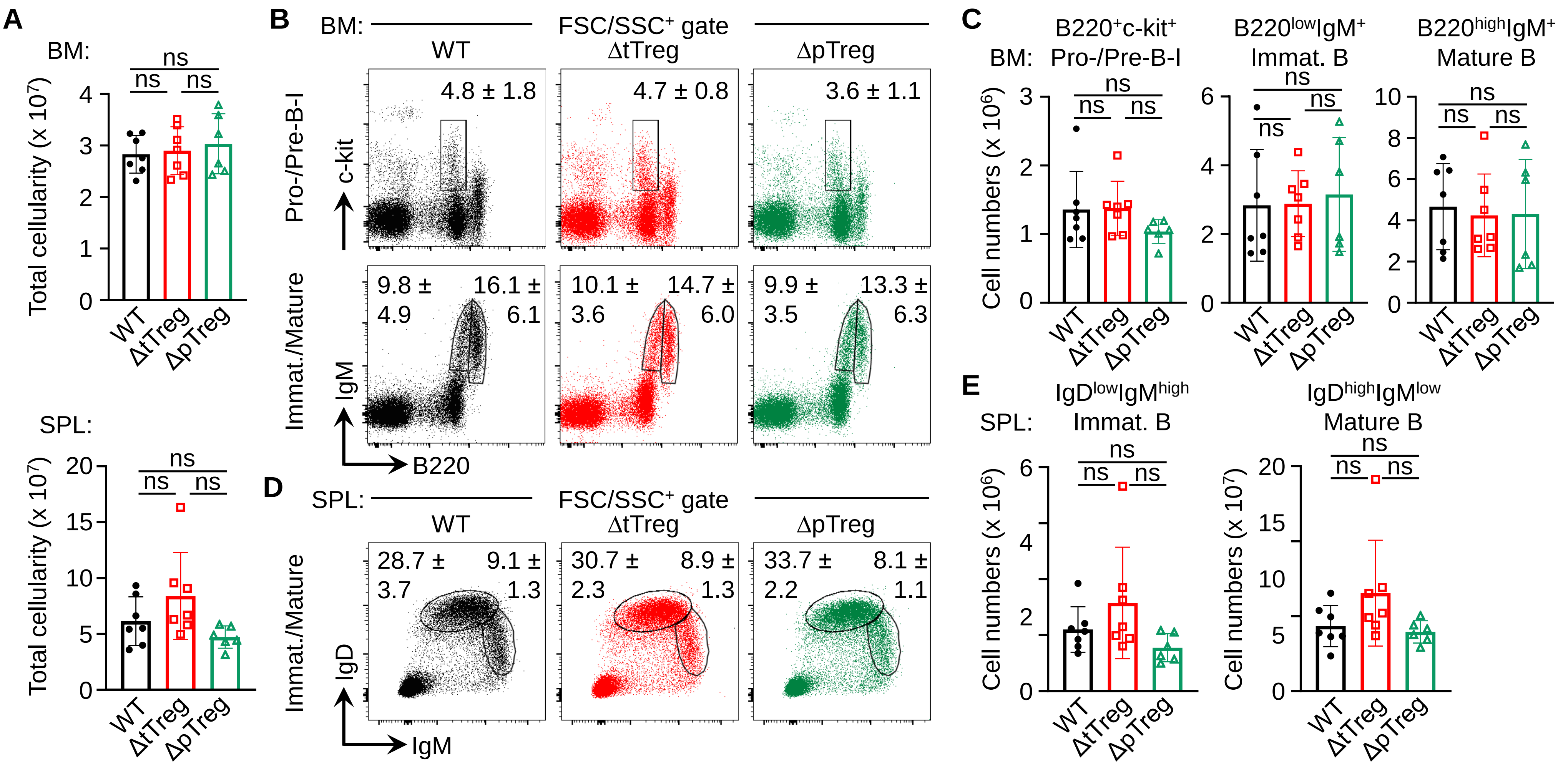

**Supplementary Figure S3: Tracking recirculating pTreg cells in the thymus of ΔtTreg mice.** Yilmazer *et al.*

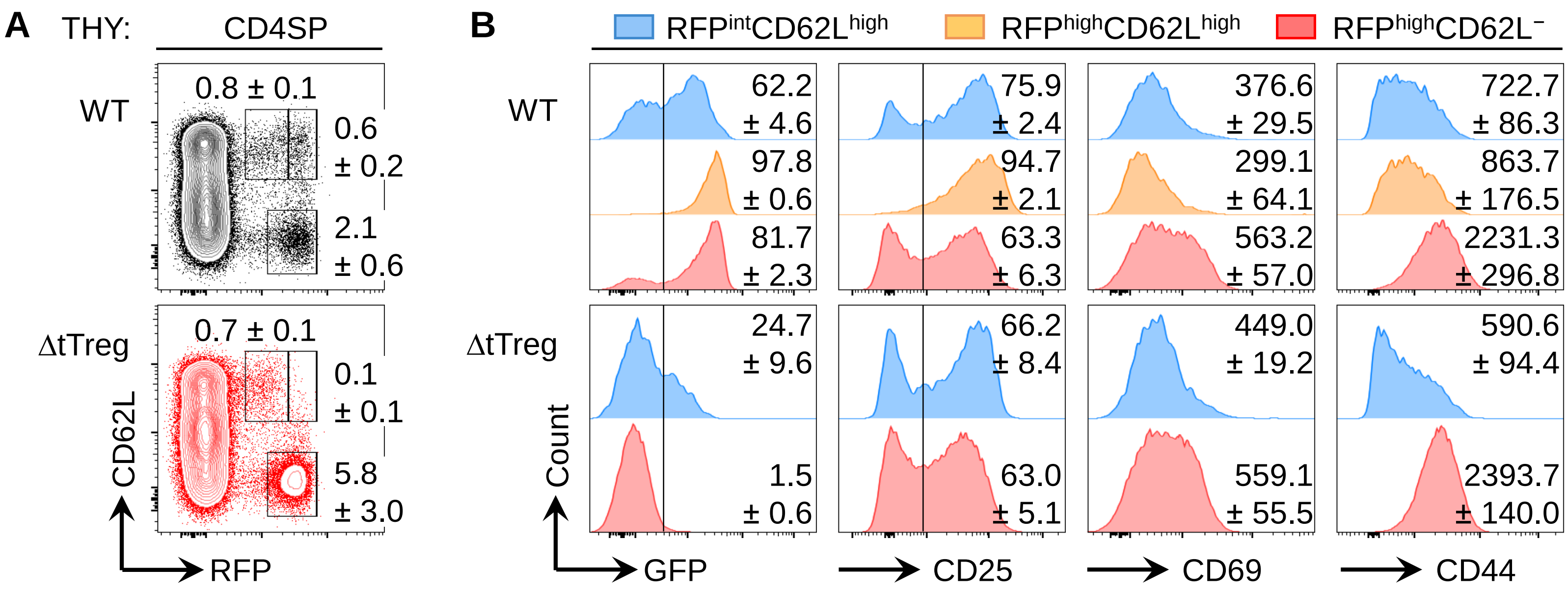

**Supplementary Figure S4: T cell numbers in peripheral lymphoid tissues of (B6>NOD)F2 ΔtTreg mice.**

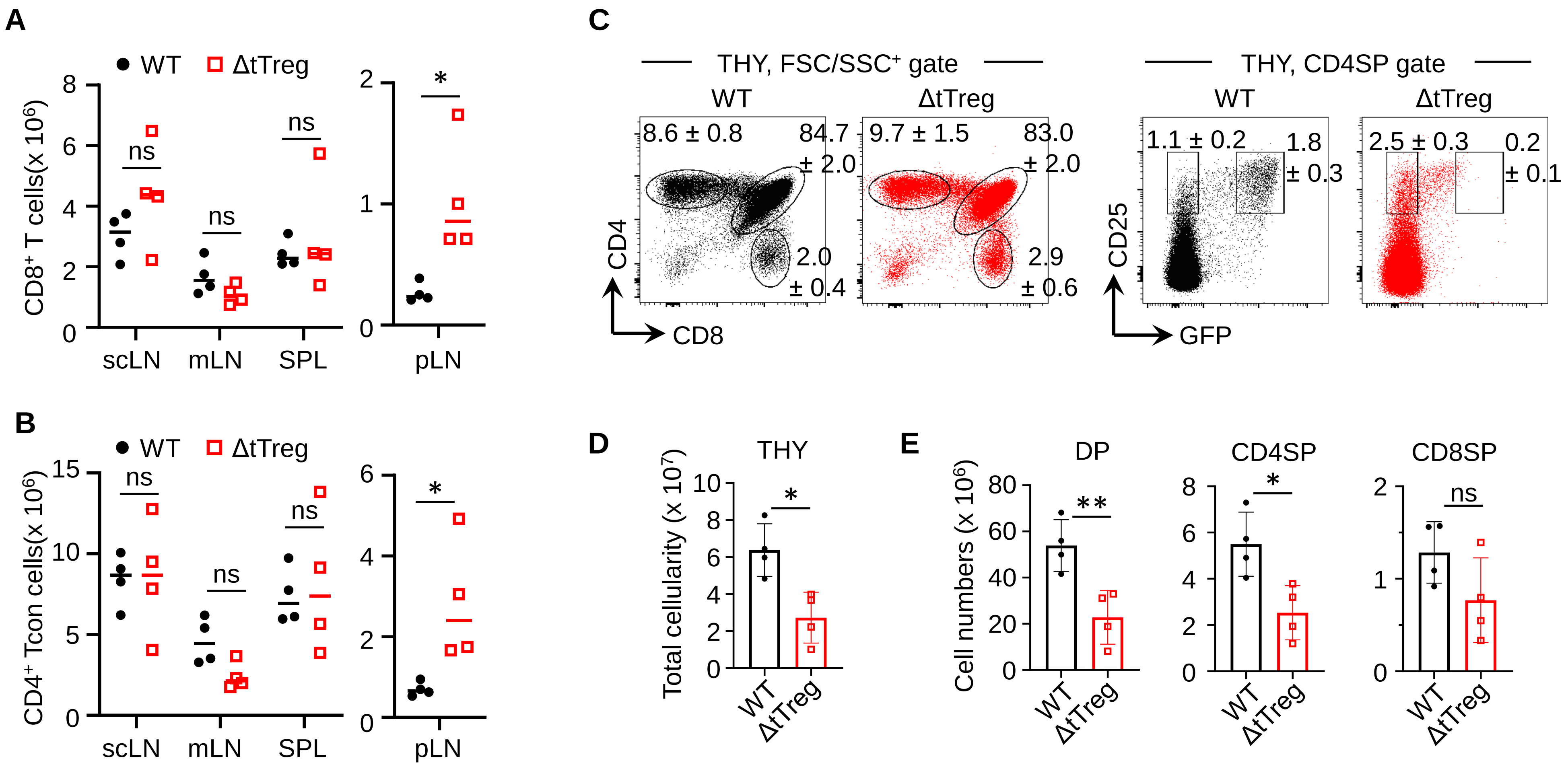

### Supplementary Figure S5: Genomic DNA-based *Idd* gene locus analysis.

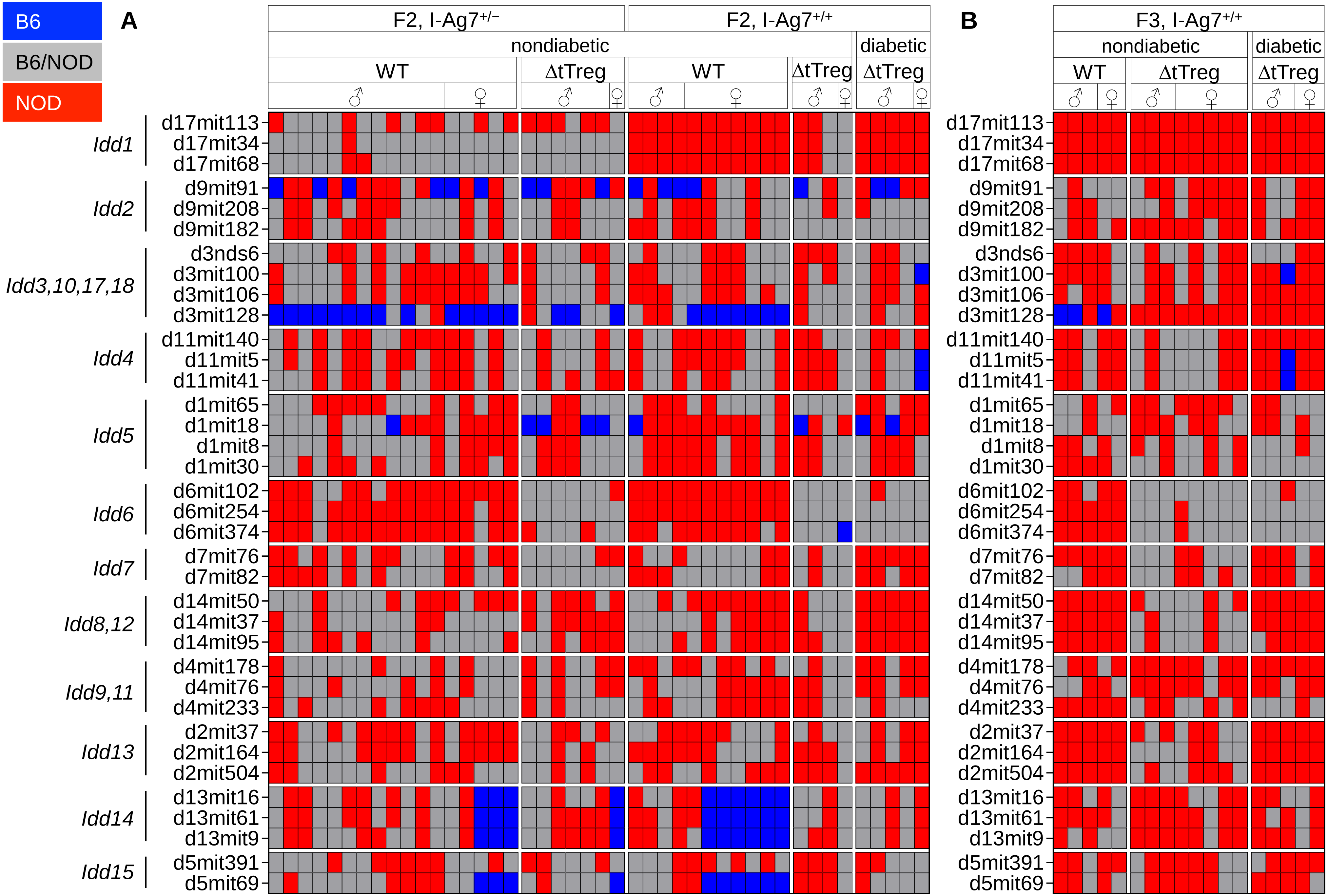
